# Comparative analysis of the molecular starvation response of Southern Ocean copepods

**DOI:** 10.1101/2023.08.17.553703

**Authors:** Cory A. Berger, Deborah K. Steinberg, Louise A. Copeman, Ann M. Tarrant

## Abstract

Large lipid-storing copepods dominate mesozooplankton biomass in the polar oceans and form a critical link between primary production and higher trophic levels. The ecological success of these species depends on their ability to survive periods of food deprivation in a highly seasonal environment, but the molecular changes that mediate starvation tolerance in these taxa are unknown. We conducted starvation experiments for two dominant Southern Ocean copepods, *Calanoides acutus* and *Calanus propinquus*, allowing us to compare the molecular starvation response between species. These species differ in life history, diet, and metabolic traits, and expressed overlapping but distinct transcriptomic responses to starvation. Most starvation-response genes were species-specific, but we identified a conserved core set of starvation-response genes related to RNA and protein metabolism. We used phylotranscriptomics to place these results in the context of copepod evolution and found that starvation-response genes are under strong purifying selection at the sequence level and stabilizing selection at the expression level, consistent with their role in mediating essential biological functions. Selection on starvation-response genes was especially strong in our focal lipid-storing lineage relative to other copepod taxa, underscoring the significance of starvation tolerance for these species. We also found that certain key lipid enzymes (elongases and desaturases) have experienced diversification and positive selection in lipid-storing lineages, reflecting the unique lipid storage needs of these animals. Our results shed light on the molecular adaptations of high-latitude zooplankton to variable food conditions, and suggest that starvation-response genes are under particularly strong sequence and expression constraints.

## 1 Introduction

The extreme seasonality of polar ecosystems causes short growing seasons for primary producers and strong seasonal variation in food availability for consumers. Even during spring and summer, food distributions in high-latitude oceans are patchy (Smetacek and Nicol, 2005; Trudnowska et al., 2016), making the ability to survive periods of food deprivation an essential trait for polar grazers. Copepods of the families Calanidae, Eucalanidae, and Rhincalanidae have evolved a conspicuous adaptation to the problem of variable food supply. These planktonic crustaceans use specialized oil sacs to sequester lipids, allowing them to survive periods of low food availability and fueling future development and reproduction (Lee et al., 2006). Copepods in these families are among the most abundant zooplankton in temperate and polar oceans and their lipid reserves can take up more than 50% of their body volume (Miller et al., 2000), making them an energy-rich food source for higher trophic levels and a critical component of productive, lipid-rich marine food webs (Record et al., 2018). Despite their ecological significance, we do not yet understand the physiological responses of these taxa to food deprivation, or the molecular and evolutionary adaptations that facilitate lipid storage and starvation tolerance.

Calanidae, Eucalanidae, and Rhincalanidae form a clade, within which oil sacs appear to be ancestral. Many species in this “oil sac clade” (OSC) undergo a period of developmentally-programmed dormancy called diapause in order to overwinter or survive other adverse conditions, usually as late-stage juveniles. Diapause is common among polar and temperate species but is not universal, and is also variable among individuals within a population (reviewed in Baumgartner and Tarrant, 2017). Species with well-defined seasonal diapause generally store unsaturated wax esters (WE) as their primary storage lipid, along with smaller amounts of triglycerides (TAG); species with lesser reliance on diapause usually have smaller lipid stores composed primarily of TAG (Lee et al., 2006). WE storage is probably ancestral in this clade and TAG storage has likely evolved independently at least twice, within *Calanus spp.* and within *Eucalanus spp.* (Fig. 1). The genetic mechanisms enabling these evolutionary transitions in lipid storage are not known and may include enzyme duplications and losses, switches in enzyme substrate preferences, and heritable changes in gene expression.

**Figure 1:**
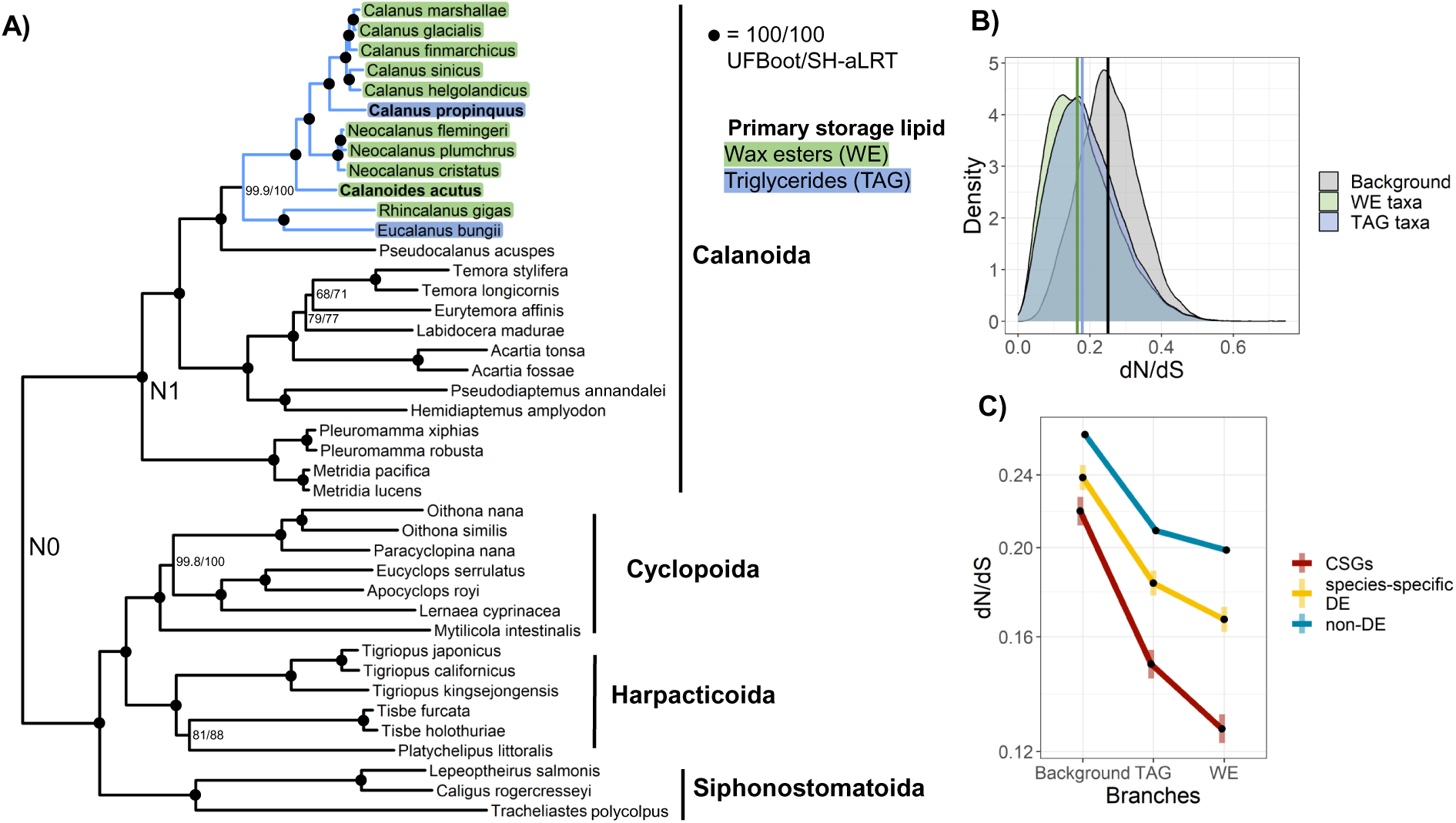
Phylotranscriptomics illuminates patterns of sequence evolution among oil sac clade copepods. A)Maximum-likelihood phylogeny of copepod species used in this study. Tree was constructed from 500 single-copy orthologs using IQ-TREE with the Q.pfam+C20+F+R5 model. Branch support values indicate ultra-fast bootstrap (UFBoot) and SH-aLRT, each calculated from 1000 replicates; black circles indicate 100% support from both metrics. Blue branches indicate the oil sac clade, and highlighted species names indicate primary storage lipid (WE or TAG). *Calanoides acutus* and *Calanus propinquus* are in bold. N0 indicates the root between Gymnoplea and Podoplea and N1 indicates the root of Calanoida; for selection analyses, we used two sets of hierarchical orthogroups inferred at these two nodes. B) Density plot of dN/dS estimates for background, WE, and TAG branches in gene trees. These estimates were derived from codeml analysis of gene trees in the N0 dataset, averaged across one-ratio, two-ratio, and three-ratio genes (see Methods). Vertical lines indicate median dN/dS for each group. C) Interaction plot of estimated marginal means of codeml dN/dS estimates from log-link Gamma GLM. Shaded bars indicate 95% confidence intervals. Note the difference in slopes; conserved starvation-response genes (CSGs) have a greater difference in dN/dS between TAG/WE and background compared to other gene sets. Y-axis is on a natural log scale.

Southern Ocean copepods exhibit diverse metabolic strategies and are less studied than their Arctic counterparts. Two of the most abundant large copepods in the Southern Ocean are *Calanoides acutus* and *Calanus propinquus*, which differ in their degree of reliance on diapause and their lipid composition. *C. acutus* performs winter diapause, stores WE, and relies primarily on a diet of large phytoplankton (Schnack-Schiel et al., 1991; Hagen et al., 1993). *C. propinquus* relies on omnivory and opportunistic feeding to overwinter, and at least part of its population remains active in surface waters during winter (Atkinson, 1991). This species primarily stores TAG and is only weakly dependent on diapause, if at all (Schnack-Schiel et al., 1991; Pasternak and Schnack-Schiel, 2002). Nonetheless, *C. acutus* and *C. propinquus* have largely overlapping circumpolar distributions and show similar spatiotemporal patterns of abundance (Marin, 1988).

The Western Antarctic Peninsula (WAP) is one of the most rapidly warming regions on Earth (Ducklow et al., 2012) and has experienced dramatic reductions in the extent and seasonal duration of sea ice cover (Stammerjohn et al., 2012). The WAP pelagic ecosystem has been studied by the Palmer Antarctica Long-Term Ecological Research program (PALLTER) since 1990, showing that regional warming is altering phytoplankton community structure and primary production (Montes-Hugo et al., 2009; Venables et al., 2013; Schofield et al., 2017; Brown et al., 2019), and zooplankton abundance and distributions (Gleiber, 2014; Steinberg et al., 2015; Thibodeau et al., 2019). *C. acutus* and *C. propinquus* both occur in the WAP, and *C. acutus* is more abundant (Marrari et al., 2011; Gleiber, 2014). Because copepods vary in their dependence on large phytoplankton, life history, and metabolic traits, responses to environmental changes will differ among species and are challenging to predict. Tolerance of short- and medium-term food deprivation will be key to copepods’ ability to acclimate and adapt to changes in an already-patchy food environment.

Although several studies have examined physiological responses of copepods to starvation (e.g Thor, 2003; Kreibich et al., 2008), there is little data on their molecular starvation response. Within Pancrustacea, insects have a well-studied starvation response involving the suppression of immune and reproductive processes and the mobilization of energy reserves through lipid and protein catabolism (McCue et al., 2017). Among copepods, field-based studies of *Neocalanus flemingeri* detected an apparent downregulation of energetic metabolism and lipid accumulation associated with low chlorophyll concentrations, suggesting some acclimatory capacity to local low food conditions (Roncalli et al., 2022). However, experiments are needed to interpret and validate field-based measures of gene expression. Studies of OSC copepods have attempted to identify molecular markers of starvation, but were limited by the measurement of only a few genes (Tarrant et al., 2021) or the use of a very short starvation period (24h) (Ohnishi et al., 2019). OSC species can survive one month or more without food even when not diapausing (Helland et al., 2003; Lee et al., 2006), so longer-term starvation experiments are required.

To address how metabolic gene expression varies across environmental conditions and experimental feeding manipulations, we sequenced and assembled transcriptomes for *C. acutus* and *C. propinquus*. In an accompanying paper, we report patterns of gene expression and other physiological metrics spanning the PAL-LTER study region (Berger et al., 2023). Here, we report ship-based starvation experiments conducted during the same research cruise. We use comparative analyses to identify conserved and divergent components of a short-term (5-9 day) starvation response, and we describe a core conserved starvation response common to both species. We then use phylotranscriptomics to infer patterns of selection associated with starvation-response genes and key lipid enzymes in the OSC. Knowledge of the copepod starvation response, including similarities and differences across species in relation to dietary, life history, and biochemical strategies, will help us understand the physiological consequences of food scarcity for these key taxa in the WAP pelagic food web.

## 2 Methods and Materials

### 2.1 Sampling and experimental design

Adult female copepods were sampled during the PAL-LTER survey cruise in austral summer 2019 aboard the ARSV *Laurence M. Gould* (detailed in Berger et al., 2023). To assess spatial variation in copepod physiology, we performed RNA-seq on *C. acutus* collected from 6 field sites and *C. propinquus* from 4 sites. See Steinberg et al. (2015) for description of PAL-LTER station grid. For starvation experiments, additional *C. acutus* were collected from site 200.040 (S67°30.664′ W70°35.377′) and *C. propinquus* from site 000.100 (S68°16.583′ W75°7.077′). Experiments were performed underway. Animals were kept in constant darkness in buckets of seawater, which was either filtered (starved; ≤3 µm filter, usually 0.45 µm) or unfiltered (fed). Buckets were surrounded with flow-through seawater to maintain temperatures similar to the sea surface. There were regular (~daily) water changes with surface seawater collected from a high-chlorophyll area as measured by the in-line ship sensor.

Animals were sampled from the experimental treatments for biochemical analyses and RNA-seq after 5 and 9 days. For sequencing, animals were stored in RNAlater at −20 °C (n=4 samples per time point). For citrate synthase activity (n=3-4 samples per time point) and lipid quantification (n=1-4 pooled samples of 2-5 individuals per time point), animals were stored at −80 °C. Samples were shipped to Woods Hole Oceanographic Institution in MA, USA on dry ice for subsequent sequencing. Lipid samples were shipped to the Marine Lipid Ecology Laboratory at the Hatfield Marine Science Center in Newport, OR, USA for analysis.

### 2.2 Photo analysis

Two independent observers scored photographs for presence/absence of food in gut and degree of egg development. Egg development was scored as 1 (no discernable eggs), 2 (moderate), or 3 (well-developed).

### 2.3 Biochemical analyses

Citrate synthase activity was determined following a modified protocol of Hawkins et al. (2016). Total lipids from individual animals were extracted and quantified using thin layer chromatography with flame ionization detection (TLC/FID) as described by Lu et al. (2008); Copeman et al. (2017), and fatty acids and alcohols were also quantified using gas chromatography. For detailed protocols, see the Supporting Information.

### 2.4 RNA extraction, library construction, and sequencing

Total RNA was extracted from individual copepods using the Aurum Fatty and Fibrous Tissue Kit (Bio-Rad) without DNase treatment and submitted to Arraystar Inc. (Rockville, MD) for library construction and sequencing. Libraries were constructed using the KAPA Stranded RNA-seq Library Preparation Kit and sequenced on an Illumina HiSeq 4000 to a depth of 40 M reads (150 bp paired-end). *C. acutus* libraries were constructed from individual samples, whereas *C. propinquus* libraries were constructed from pooled equivalent amounts of RNA from two individuals. Sequence quality was assessed with FASTQC v0.11.7 (https://www.bioinformatics.babraham.ac.uk/projects/fastqc/).

### 2.5 Quality filtering and transcriptome assembly

For *C. acutus*, we used an individual-based approach in which six individual transcriptomes were assembled and then merged. Sequences were error-corrected using RCorrector v1.0.4 (Song and Florea, 2015) and adapter and quality trimmed using TrimGalore v0.6 (https://github.com/FelixKrueger/TrimGalore) with settings “–quality 5 –length 50.” Ribosomal RNA was removed with SortMeRNA v2.1b (Kopylova et al., 2012). Six individual libraries were assembled using Trinity v2.9.1 (Grabherr et al., 2011) with default parameters, and the resulting assemblies were combined using TransFuse v0.5.0 (https://github.com/cboursnell/transfuse). Since *C. propinquus* libraries consisted of pooled individuals, we used a standard Trinity pipeline in which reads from eight libraries were concatenated prior to assembly. Sequences were error-corrected using RCorrector and trimmed with TrimGalore v0.6 with default settings. A transcriptome was then assembled *de novo* using Trinity v2.6.6. Accessions for reads used in transcriptome assemblies are in Table S1.

Assemblies were clustered using CD-HIT-EST v4.7 (Li and Godzik, 2006) at 98% sequence similarity (-c 0.98 -b 3). We predicted peptide sequences with TransDecoder v5.5.0 (https://github.com/TransDecoder/TransDecoder), and also retained peptides with hits (e-value = 1 × 10^−10^) to Swiss-Prot (Boutet et al., 2007, March 18, 2020) searched by DIAMOND BLASTP v0.9.30 (“more-sensitive” mode; Buchfink et al., 2015) or PFAM (Finn et al., 2014, March 16, 2020) searched by hmmscan (http://hmmer.org). Assemblies were annotated with Trinotate (Bryant et al., 2017). Sequences for which the top five BLASTP hits by e-value contained only non-metazoan sequences were considered contamination and removed. Transcriptome completeness was assessed with BUSCO v4 (Arthropoda odb10 database; Simão et al., 2015).

### 2.6 Expression analyses

We quantified transcript abundance with Salmon v1.1.0 (Patro et al., 2017) with “-gcBias-seqBias -validateMappings”. Transcripts were clustered using Corset v1.09 (Davidson and Oshlack, 2014) and counts were summarized to gene level using tximport v1.20.1 (Soneson et al., 2016) with “countsFromAbundance = lengthScaledTPM”. We retained genes with at least 15 counts in 4 libraries. Based on initial visualization and clustering, we removed one outlier library in each species (Figs. S1 & S2). We conducted principal component analysis (PCA) with the “prcomp” package. For differential expression (DE) analysis, we applied limma-voom (Law et al., 2014) with sample-specific quality weights and robust empirical Bayes moderation, using the linear model “ 0 + group + food*day + site”. We considered genes DE at FDR<0.05. For discriminant analysis of principal components, we used the “adegenet” package v2.1.8 (Jombart, 2008). Gene ontology (GO) enrichment was performed with GO MWU (Wright et al., 2015) with genes ranked by log2 fold-change (LFC). We used REVIGO (Supek et al., 2011) to visualize and summarize significant GO terms. Significance of correlations between orthologs was assessed with permutation tests in which orthology relationships were shuffled (1 × 10^5^ permutations). Weighted Gene Co-expression Network Analysis was performed using the “WGCNA” R package (Langfelder and Horvath, 2008). For each species, we constructed one network from all experimental and field samples using parameters corType = “bicor”, maxPOutliers = 0.2, power = 7, networkType = “signed hybrid”, TOMType = “signed”, minModuleSize = 30, and mergeCutHeight = 0.25. Eigengenes were associated with treatment conditions using the same linear model as the DE analysis.

To identify genes with similar fold-change across species, we used an equivalence test based on the “Two One-Sided Tests” procedure (Schuirmann, 1987). We performed this test on ortholog pairs with at least one DE gene in either species. We considered LFCs as “equivalent” if the LFC of the non-DE ortholog was within ± 50% of the DE ortholog, or if the DE gene with the smaller LFC was within ± 50% of the DE gene with the larger LFC. We obtained moderated standard errors and degrees of freedom for LFC estimates from limma and calculated pooled degrees of freedom for ortholog pairs with the Welch-Satterhwaite equation (Satterthwaite, 1946). The equivalence test p-value is the greater of the two p-values obtained by left- and right-tailed Welch’s t-tests (Welch, 1947). P-values were corrected for multiple testing using the Benjamini-Hochberg procedure. We extended this approach to multi-copy orthogroups by testing for equivalence between the weighted means of LFC estimates within each orthogroup across species, with weights corresponding to the inverse squared standard errors of LFC estimates. Once again, we performed this test on orthogroups with at least one DE gene in either species. As a metric for expression divergence, we computed the inverse of the above equivalence test, thus obtaining a p-value for the probability that the difference in LFC was greater than 50%.

We estimated dispersion of gene expression from field samples as in Fair et al. (2020). This assumes that gene expression follows a Gamma distribution and that variability due to the measurement process follows a Poisson distribution. We fit negative binomial regressions to each gene to estimate Poisson overdispersion, providing an estimate of biological variability. We then regressed out mean expression, resulting in a measure of dispersion relative to the variability expected given a gene’s expression level. We tested for differences in dispersion between groups of genes with a weighted linear model (weights were inverse squared standard error of dispersion estimates).

### 2.7 Phylotranscriptomics

We obtained or re-assembled transcriptomes of 39 other copepod species using published data (assembly details in Table S1). We assembled new transcriptomes using a common trimming and assembly pipeline in cases where only raw reads were available or existing assemblies had low BUSCO scores (full methods in Supplemental Information). We ran OrthoFinder v2.5.2 (Emms and Kelly, 2015) with “-m MSA” on this dataset of 41 protein-coding transcriptomes to infer gene families. We constructed maximum-likelihood species trees from a set of 500 single-copy genes using an edge-proportional partition model and a site-heterogeneous posterior mean site frequency model (Q.pfam+C20+F+R5 Wang et al., 2018) implemented in IQ-TREE. These genes were processed to mitigate potential sources of bias and long-branch attraction (full methods in Supplemental Information), and both trees recovered identical topologies (Fig. 1A); Fig. S3).

#### 2.7.1 Selection analyses

We analyzed selection in two datasets: 1) orthogroups present in at least 21/41 species and within the OSC (N0 dataset; n=10,487); and 2) orthogroups inferred within Calanoida and present in at least 13/25 calanoid species (N1 dataset; n=13,640). For each orthogroup, we aligned peptide sequences with MAFFT L-INS-I and created codon alignments with Pal2nal v14.1 (Suyama et al., 2006). We removed codons with >90% gaps and constructed trees using FastTree v2.1.10 (Price et al., 2009) with parameters “-lg -gamma -pseudo”. We then re-optimized branch lengths using RaxML-NG v1.0.1 (Stamatakis, 2014) with the LG+FO+I+R4 model, and removed long branches using TreeShrink (alpha=0.05).

We estimated branch-specific rates of sequence evolution (dN/dS) with the MG94 codon model using FitMG94 (https://github.com/veg/hyphy-analyses/tree/master/FitMG94) in Hyphy (Pond et al., 2005). We tested for differences in dN/dS between groups of genes with a weighted generalized linear model accounting for expression level (GLM with Gamma error and log link; weights were inverse squared standard error of dN/dS estimates). To identify genes with different selection regimes between the OSC (foreground) and other lineages (background), we compared branch models using codeml (Yang, 2007). For each gene, we fit three models: 1) a one-ratio model with one dN/dS ratio (ω) for the entire tree, 2) a two-ratio model with separate ω for OSC and background lineages, and 3) a two-ratio model with OSC ω fixed at 1. We ran analyses with two starting values of ω (0.5 and 2) and considered runs to converge if ω estimates were within ±0.01 across both starting values; this was the case for 8613/10,487 (82%) N0 and 11,645/13,640 (85%) N1 orthogroups. All models were run with parameters “CodonFreq = 2, fix kappa = 0, kappa = 1, and fix blength = 2”. For genes best fit by the two-ratio model (p<0.05), we additionally fit a model in which TAG-storing species (*C. propinquus* and *Eucalanus bungii*) had a third ω. We tested for differences in ω estimates between groups of genes and lineages with an unweighted GLM (Gamma error and log link). For genes with higher ω in the foreground, we used two follow-up tests within Hyphy, BUSTED-PH (https://github.com/veg/hyphy-analyses/tree/master/BUSTED-PH) and RELAX (Wertheim et al., 2015). Interactions were retrieved from GLMs with the “emmeans” package (Lenth et al., 2023).

To annotate fatty acid elongase (ELOV) and desaturase (FAD) sequences, we down-loaded representative metazoan peptide sequences from NCBI (accessions in Table S2), built profile alignments with MAFFT L-INS-I and hmmer v3.3.1 (Eddy, 2011), and used hmm-search (e-value < 1 × 10^−10^) to identify candidate peptides. We used hmmscan to search PFAM (downloaded March 15, 2020) and retained sequences with hits (e-value < 1 × 10^−3^) to representative domains (“ELO” for ELOV, “FA desaturase” for FAD) and a minimum length of 100 amino acids. We also required ELOV sequences to have at least 5 transmembrane domains predicted by TMHMM v2.0 (Krogh et al., 2001) and a histidine box motif (HxxHH/QxxHH). We used BLASTP against the nr database to check sequences for correct taxonomic assignment and removed contaminant sequences. Sequences passing these checks were clustered at 85% identity within species using CD-HIT (-c 0.85 -G 0 -aS 0.7). We aligned sequences using MAFFT L-INS-I, trimmed sites with >90% gaps, and constructed gene trees with IQ-TREE after model selection with ModelFinder. We used Pal2nal to construct codon alignments. The ELOV family contains deep paralogs with two ancestral copies in Metazoa (Hashimoto et al., 2008), so these paralogs were analyzed separately; we named these clades based on homology to vertebrate ELOVs, which are numbered 1-7 (see Supplement for full trees). Within the ELOV1/4/7 clade, we recovered a clade of putative elongases present in 8 harpacticoid and cyclopoid species, on a long branch sister to the remaining copepod ELOV1/4/7 sequences. We dropped this clade from our analysis because it contained no foreground sequences and may be an out-paralog. We used RELAX and BUSTED-PH as above to test for differences in selection between foreground and background.

## 3 Results

### 3.1 Transcriptome assembly, species tree, and homology inference

We assembled transcriptomes for *Calanoides acutus* (Bioproject PRJNA757455) and *Calanus propinquus* (Bioproject PRJNA669816). The assemblies contained 73,545 and 102,391 protein-coding transcripts, respectively, with BUSCO scores of 96.6% and 85.5%, respectively. We also obtained or re-assembled transcriptomes of 39 other copepod species (Table S1). Together, these 41 transcriptomes span the major copepod orders (Calanoida, Harpacticoida, Cyclopoida, and Siphonostomatoida), with 12 species belonging to the oil sac clade (OSC). We used this dataset to infer gene families and construct a maximum-likelihood species tree, which recovered major clades of copepods with strong support (Fig. 1A). Using this phylogeny, OrthoFinder inferred hierarchical orthogroups, including 9087 one-to-one orthologs and 37,270 multi-copy homologs between *C. acutus* and *C. propinquus*.

### 3.2 Differential gene expression

We conducted shipboard starvation experiments with field-collected *C. acutus* and *C. propinquus*; we performed RNA-seq on fed and starved animals collected after 5 and 9 days, and animals collected directly from various field sites. For both species, principal component analysis (PCA) clearly separated experimental and field samples, with some separation of fed and starved animals but no clustering by time point (Fig. 2). Only one gene in either species had a significant interaction between starvation and time, indicating that gene expression changes were largely concordant between time points (Tables 1, 2). We therefore focus on comparisons between all starved and fed samples. We identified 3687 (6.2% of transcriptome) and 1185 (1.5%) DE genes between starved and fed animals in *C. acutus* and *C. propinquus*, respectively (FDR<0.05). Because *C. acutus* were more abundant during sampling, we included a “refeeding” group in which animals were starved until day 5 and refed until day 9. There were no DE genes between refed and fed animals, and discriminant analysis of principal components confirmed that refeeding mostly reversed gene expression changes between fed and starved animals (Fig. 2C).

**Figure 2:**
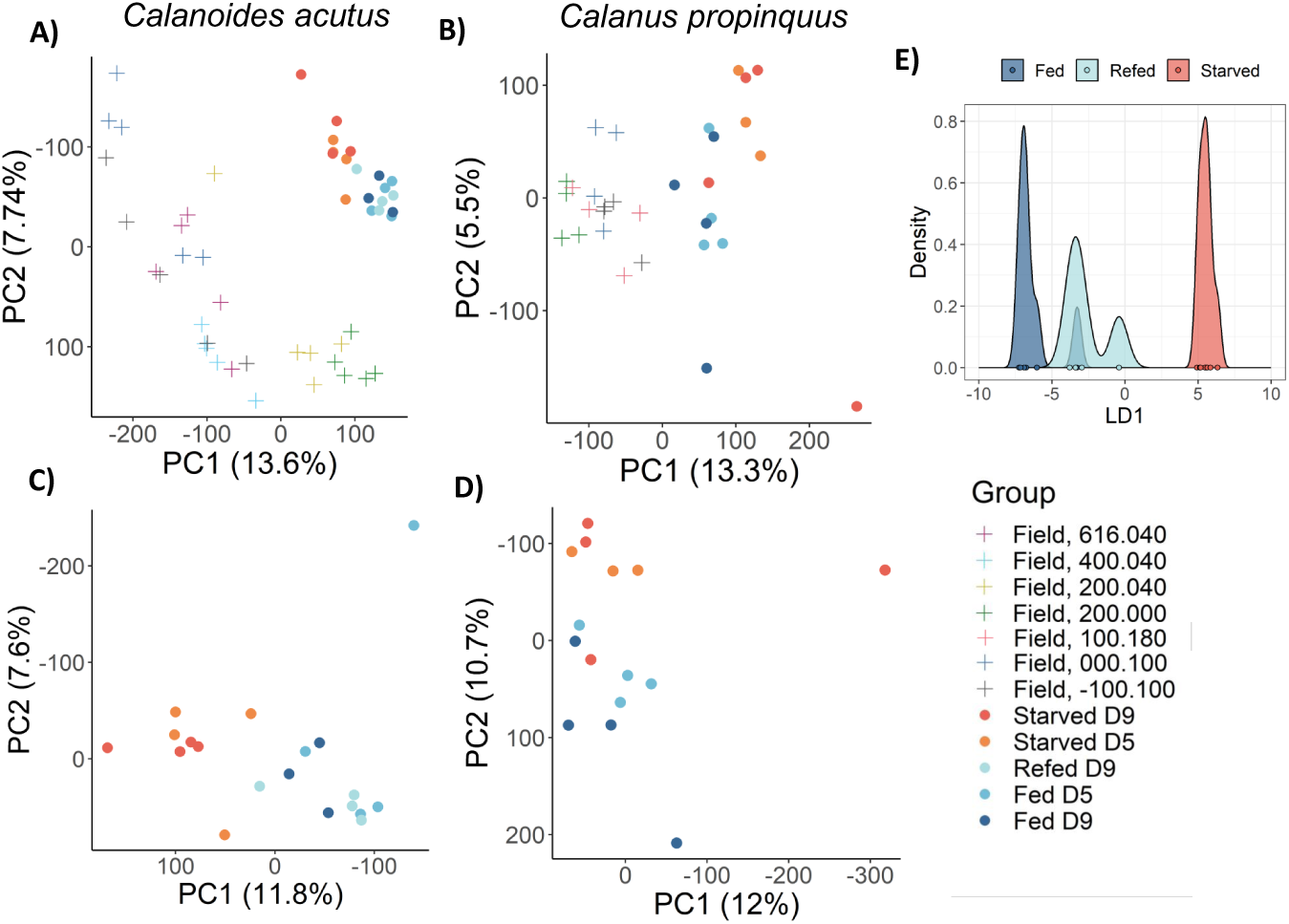
Principal component analysis (PCA) separates field, fed, and starved samples in *Calanoides acutus* (left) and *Calanus propinquus* (middle). A) and B) show PCAs calculated using all samples; C) and D) were calculated using only experimental samples. PCA was performed on log-scaled counts-per-million of all genes expressed above our expression cutoff. Percentages indicate variance explained by each PC. Coordinates correspond to the PAL-LTER sampling grid. Experimental *C. acutus* were collected from site 200.040, and *C. propinquus* from site 000.100. E) Discriminant analysis of principal components showing refed *C. acutus* overlap with fed samples. LD1 is the discriminant axis that maximizes the variation between fed and starved samples.

**Table 1:**
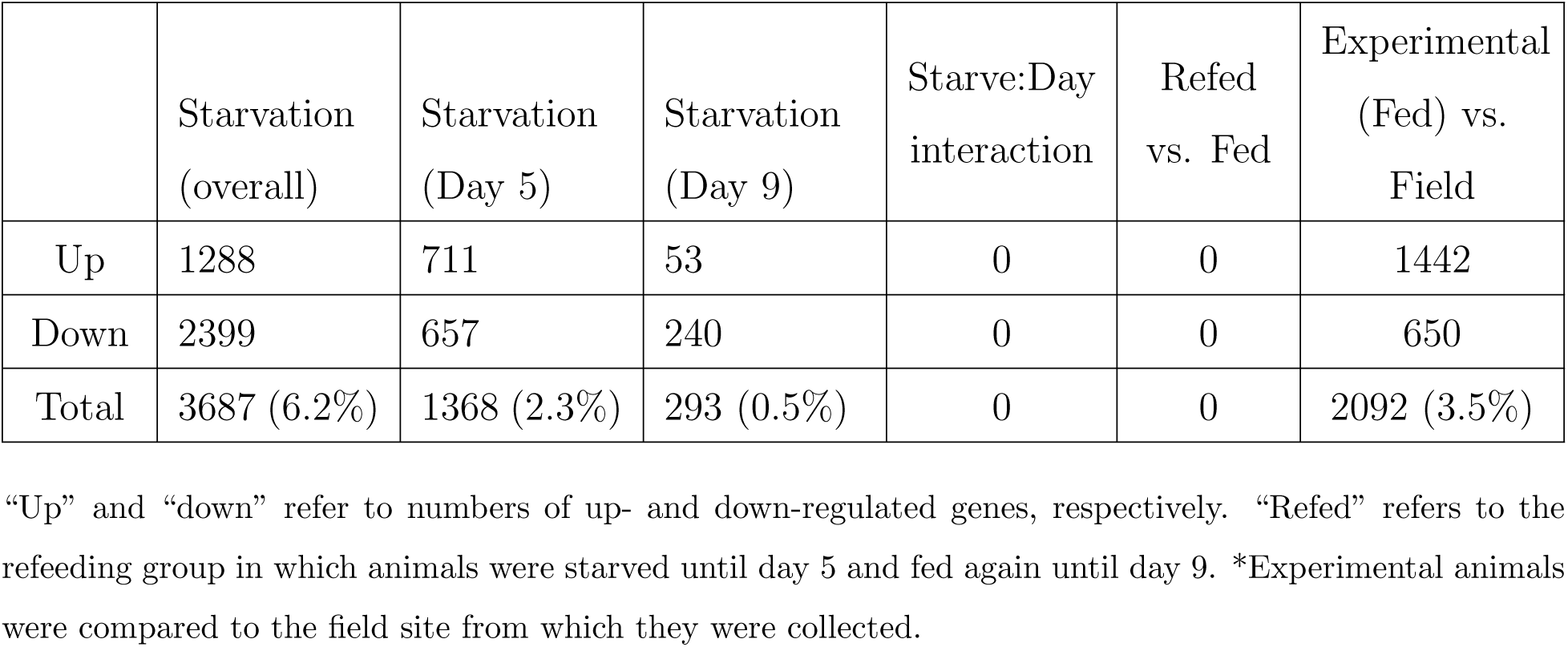
Numbers of differentially expressed genes in the copepod *Calanoides acutus*.

|  | Starvation<br>(overall) | Starvation<br>(Day 5) | Starvation<br>(Day 9) | Starve:Day<br>interaction | Refed<br>vs. Fed | Experimental<br>(Fed) vs.<br>Field |
| --- | --- | --- | --- | --- | --- | --- |
| Up | 1288 | 711 | 53 | 0 | 0 | 1442 |
| Down | 2399 | 657 | 240 | 0 | 0 | 650 |
| Total | 3687 (6.2%) | 1368 (2.3%) | 293 (0.5%) | 0 | 0 | 2092 (3.5%) |

“Up” and “down” refer to numbers of up- and down-regulated genes, respectively. “Refed” refers to the refeeding group in which animals were starved until day 5 and fed again until day 9. \*Experimental animals were compared to the field site from which they were collected.

**Table 2:** Numbers of differentially expressed genes in the copepod *Calanus propinquus*.

|  | Starvation<br>(overall) | Starvation<br>(Day 5) | Starvation<br>(Day 9) | Starve:Day<br>interaction | Experimental<br>(Fed) vs.<br>Field |
| --- | --- | --- | --- | --- | --- |
| Up | 687 | 0 | 142 | 0 | 1173 |
| Down | 498 | 0 | 132 | 1 | 674 |
| Total | 1185 (1.5%) | 0 | 274 (0.4%) | 1 | 1847 (2.4%) |

“Up” and “down” refer to numbers of up- and down-regulated genes, respectively. Experimental animals were compared to the field site from which they were collected.

### 3.3 Physiological assays

We assessed gut content and reproductive condition from photo analysis. In both species, some individuals in the “starved” treatment had visible food in gut, although much less than “fed” treatments (see Discussion and Fig. S4). There were no significant differences in reproductive status for either species. No *C. propinquus* died during the experiment, but mortality of *C. acutus* increased from 2-8% by day 5 to 20-36% by day 9, with no difference between fed and starved groups (χ^2^, p=0.39). Starved animals of both species had lower citrate synthase activity than fed animals (Fig. 3A; ANOVA, p<0.05).

**Figure 3:**
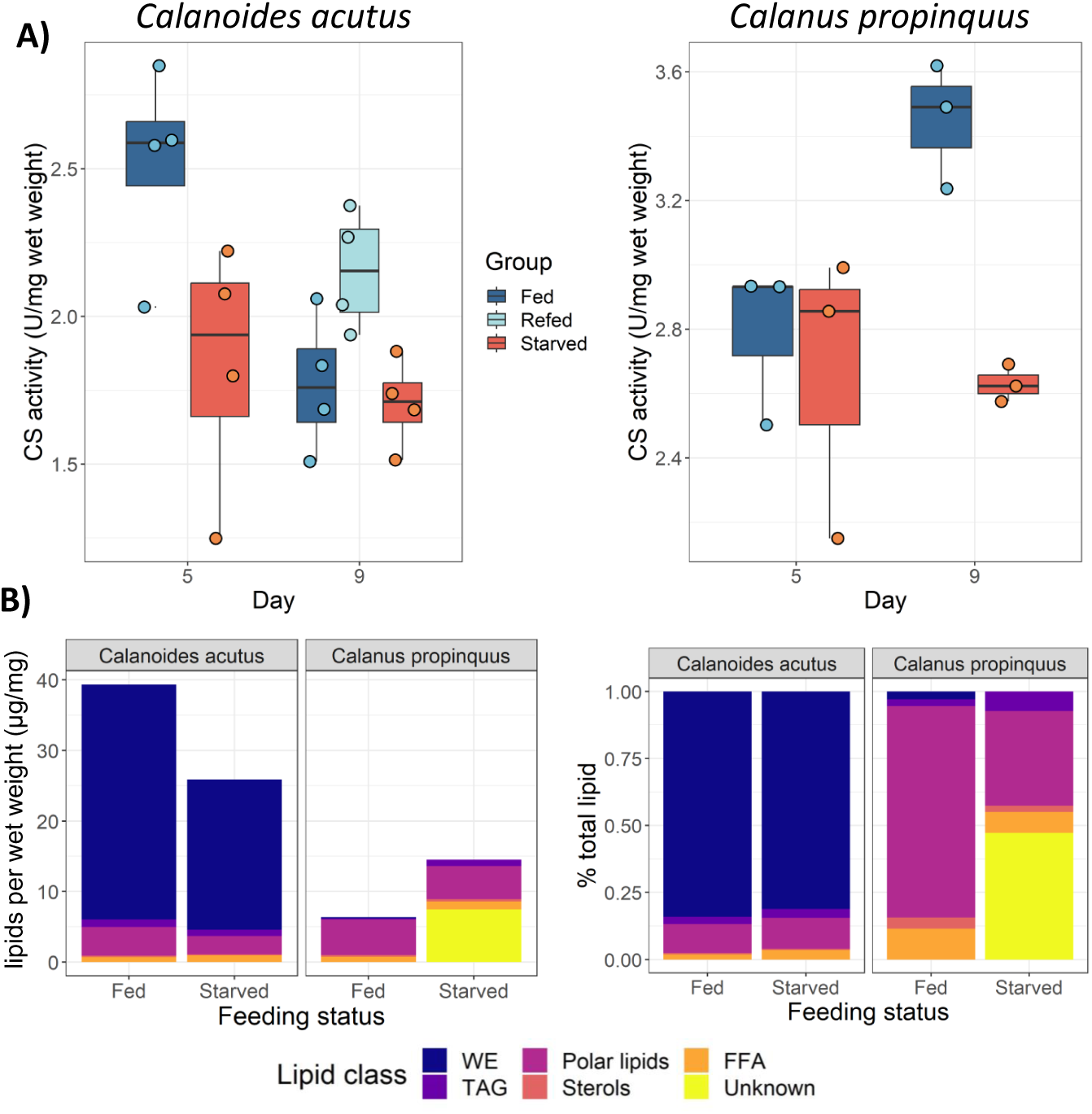
Effects of starvation on citrate synthase activity and lipid content. A) Enzyme activity normalized to wet weight in *C. acutus* (left) and *C. propinquus* (right); in both species, starved animals had lower CS activity than fed (ANOVA, p<0.05). B) Lipid classes per wet weight (µg mg^−1^; left) and as a proportion of total lipids (right).

We quantified total lipids, lipid classes, and fatty acid and alcohol (FA+ALC) composition (mass percentage). Lipids were assigned to standard lipid classes—wax esters (WE), triglycerides (TAG), sterols, polar lipids, and free fatty acids (FFA). We also detected an unknown neutral lipid in three *C. propinquus* samples; data were analyzed with and without this lipid, and its inclusion did not affect the results. Starved *C. acutus* had lower total lipid content than fed animals, with mean 86±34 vs. 147±37 µg per individual (Fisher-Pitman test, p=0.038, n=10; Fig. 3B). Starved *C. propinquus* did not differ in total lipid content (p=0.5, n=5; Fig. 3B). There was no significant difference in any particular lipid class between starved and fed animals for either species (p>0.05). Using correspondence analysis (CA), fed and starved *C. acutus* did not differ in FA+ALC composition (CA constrained by feeding status, ANOVA, p=0.4). For *C. propinquus*, CA constrained by feeding status was significant (ANOVA, p=0.017), but our sample size was small (n=3 fed, n=2 starved). In summary, there were not clear differences in lipid or FA+ALC composition between fed and starved animals of either species, although starved *C. acutus* had reduced lipid content overall.

### 3.4 Species share a core starvation response regulating RNA and protein metabolism

*C. acutus* and *C. propinquus* showed broadly similar transcriptomic responses to starvation (full DE and GO enrichment results in Supplemental File 2). The log2 fold-change (LFC) between starved and fed samples was correlated among one-to-one orthologs (Spearman’s ρ=0.32), as was the weighted average LFC of multi-copy homologs (ρ=0.34; Fig. 4A). These correlations, and all others discussed in this paper, were highly significant as assessed by permutation tests (p<1 × 10^−5^). Gene ontology (GO) enrichment of starvation-response genes revealed functional similarities between species. In both species, upregulated genes were enriched for terms related to RNA and protein metabolism, the cell cycle, DNA repair, and response to stress, while downregulated genes were enriched for terms related to development, transposition, transcription factor activity, chromosome organization, and sulfur compound metabolism.

**Figure 4:**
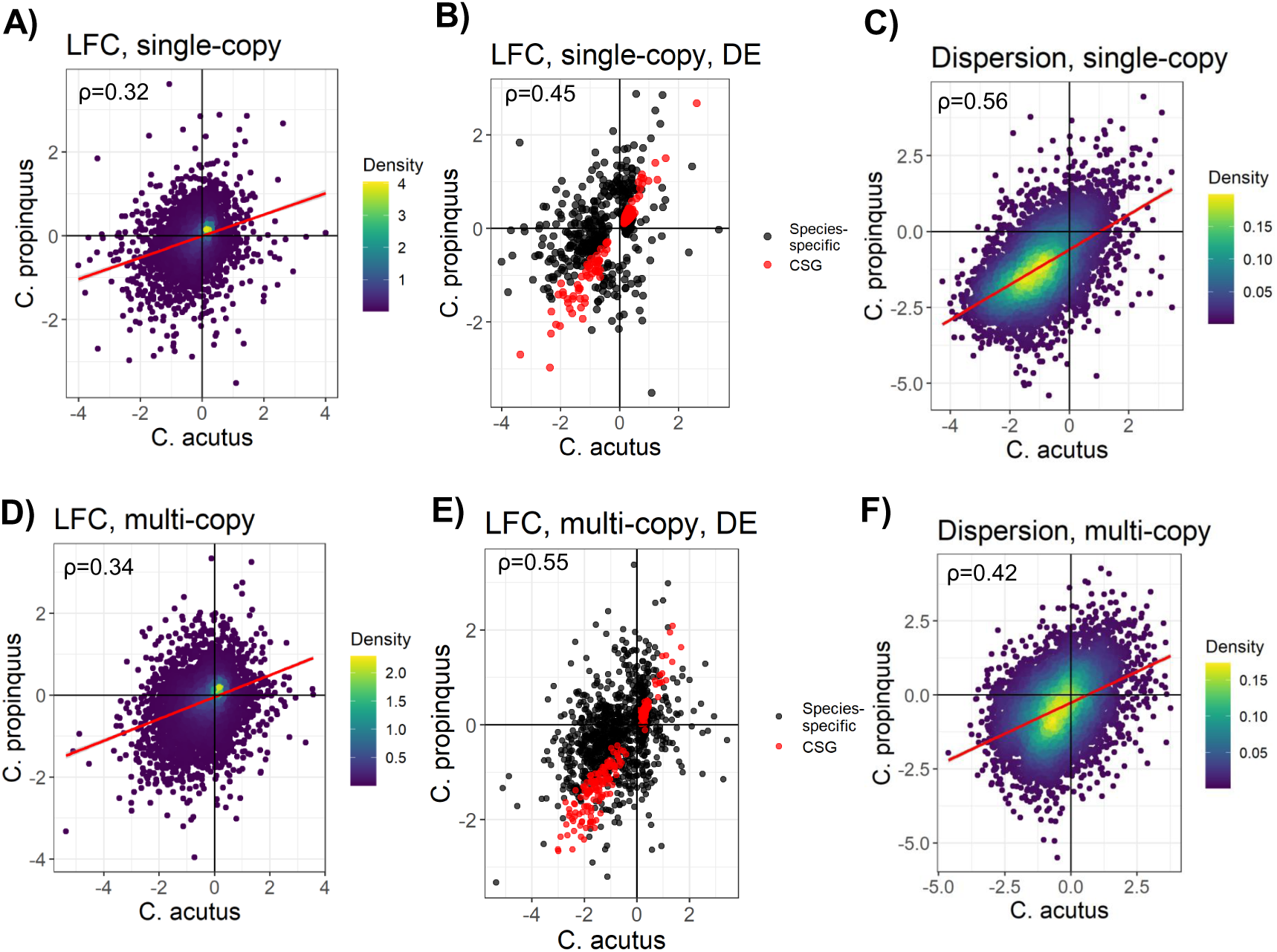
Gene expression responses to starvation are broadly correlated across copepod species. Correlation of starvation LFC for single-copy genes (A); single-copy genes that were DE in at least one species (B); multicopy genes (D); and multi-copy genes that were DE in at least one species (E). Correlation of dispersion estimates calculated using field samples for single- (C) and multi-copy (F) genes. For multi-copy genes (bottom row), points represent the average value of the gene family weighted by the inverse squared standard error of each gene’s estimate. Conserved starvation-response genes (CSGs) are in red.

In *C. acutus*, weighted gene co-expression network analysis WGCNA identified 6 modules associated with starvation (Fig. S5; File S3). Three modules containing 6685 genes had higher expression in starved animals, and were enriched for GO terms related to RNA metabolism, protein ubiquitination, and regulation of cell cycle processes (Black module); DNA replication and repair, stress response, protein localization and transport, tRNA metabolism, and macromolecule biosynthesis and catabolism (Magenta); and translation, protein folding, ribosomes, proteasomes, and mitochondrial respiration (Yellow). Three modules containing 15,608 genes had lower expression in starved animals. These were enriched for GO terms related to lipid biosynthesis, fatty acid metabolism, nucleotide biosynthesis, ion transport, and sulfur compound and carbohydrate metabolism (Red); oxidoreductase activity, amino acid metabolism, carnitine biosynthesis, mitochondrial transport, and organic acid transport (Salmon); and development and cell proliferation, reproduction, immune response, and peptidase and hydrolase activity (Turquoise).

In *C. propinquus*, WGCNA identified 8 modules associated with starvation (Fig. S6; File S4). Five modules containing 3312 genes had higher expression in starved animals, and were enriched for GO terms related to protein catabolism and autophagy (Darkmagenta); amino acid and vitamin metabolism (Darkorange); RNA and protein transport, the cell cycle, translation, DNA metabolism and repair, and stress response (Magenta); and response to insulin and peptide signalling, triglyceride metabolism and biosynthesis, lipid catabolism, and carboxylic acid catabolism (Purple)–the remaining module was not enriched for any GO terms. Three modules containing 19,440 genes had lower expression in starved animals. These were enriched for GO terms related to phagocytosis and immune response (Paleturquoise); oxidoreductase activity, carboxylic acid, amino acid, and carbohydrate metabolism (Royalblue); and fatty acid metabolism, DNA replication, development, and cell proliferation (Turquoise).

To define a conserved starvation response, we used an equivalence test to identify singlecopy orthologs with LFC within ±50% of one another, conditioned on one ortholog being DE in either species (see Methods). We identified 337 pairs of conserved starvation-response orthologs (FDR<0.05; Fig. 4B). We extended this analysis to multi-copy gene families by comparing the weighted average LFC within each orthogroup and species. We identified 223 multi-copy orthogroups with equivalent LFC (FDR<0.05; Fig. 4E), containing 1232 genes. Although this method does not require all genes within an orthogroup to respond in the same way, just 7/1232 of these genes had an LFC that differed in sign and differed from the group mean (One-tailed t-test, p<0.05, Fig. S7), indicating that in practice nearly all genes within these orthogroups had similar LFCs. Overall, single- and multi-copy “conserved starvation-response genes” (CSGs) represented 14.5% of all DE genes in both species (Fig. 4B). Upregulated CSGs were enriched for GO terms related to RNA and protein metabolism (e.g. transcription, splicing, translation, ribosome biogenesis, nuclear export, and protein catabolism), and DNA repair and replication; downregulated CSGs were enriched for DNA-mediated transposition and DNA polymerase activity. Thus, although most starvation-induced gene expression changes were species-specific, we identified a conserved set of starvation-response genes that mediate fundamental aspects of cellular metabolism.

We specifically analyzed genes annotated with any descendent GO terms of “lipid metabolism” (GO:0006629), “protein metabolism” (GO:0019538), and “response to stress” (GO:0006950). Several terms enriched among CSGs were related to protein metabolism (ubiquitination and regulation of translation) and stress (DNA repair), but none were related to lipid metabolism. Protein- and stress-related genes were enriched among CSGs compared to species-specific DE genes (Wilcox test, p<0.005), but this was not true for lipid-related genes (p=0.06), suggesting that protein metabolism and stress response genes were relatively conserved in their starvation response, while lipid metabolism genes were divergent. To test this, we computed a gene-wise metric of similarity for the starvation response corresponding to the probability that the difference in LFC between species was greater than 50% (the inverse of the above equivalence test). Expression of lipid-related genes was more divergent compared to protein-related, stress-related, and all non-lipid related genes (Wilcox, p<0.05). Protein-related genes, meanwhile, had more similar expression profiles across species than non-protein related genes, lipid-related genes, and stress-related genes (Wilcox, p<0.05). Therefore, genes that mediate protein metabolism generally responded similarly to starvation in *C. acutus* and *C. propinquus*, while lipid metabolism genes generally differed in their response.

That said, some key enzymes with known roles in lipid metabolism were CSGs. Both species upregulated a long-chain-fatty-acyl-CoA ligase, a family of CREB-like genes, and the TOR kinase, and downregulated fatty acid desaturases (FADs), fatty acid elongases (ELOVs), and various hydrolases and esterases. Several species-specific DE genes also had roles in lipid metabolism, including additional elongases, a lipase, cholesterol transporters, and acyl-CoA dehydrogenases and reductases (*C. acutus*); and various lipases, desaturases, and lipoprotein receptors (*C. propinquus*). Both species modulated genes related to insulin/TOR signalling: *C. propinquus* upregulated FoxO, an insulin-like peptide, an insulin receptor, and several insulin-degrading enzymes, and *C. acutus* upregulated homologs of CREB and Myc. Additionally, both species strongly downregulated vitellogenins and upregulated a homolog of the ecdysone receptor.

#### 3.4.1 Comparison with an experimental handling response

We noted large differences in gene expression between experimental and field samples (2092 and 1847 DE genes in *C. acutus* and *C. propinquus*, respectively; Fig. 2), which we interpret as a generalized stress response to handling and captivity. In both species, these genes were enriched for GO terms related to stress and immune response, muscle development, and cytoskeleton organization. Using the same equivalence test used to define CSGs, only 7.2% (284/3939) of handling-response genes had equivalent expression across species (Fig. S8), compared to 14.5% of starvation-response genes (709/4872). The correlation between homologous genes for the handling response was also weaker than for the starvation response (Fig. S8; ρ=0.27 for single-copy, ρ=0.15 for multi-copy). Thus, the starvation response was more similar between species than the generalized response to captivity, and should be interpreted as occurring on top of this pre-existing handling stress.

#### 3.4.2 Starvation-response genes have high expression and low dispersion

Starvation-response genes were more highly-expressed than non-DE genes in both species, and CSGs had higher expression than species-specific starvation-response genes in *C. acutus* (LM, p<1 × 10^−4^). To study the expression properties of starvation-response genes, we estimated mean-corrected dispersion of gene expression across field samples—thus avoiding variance introduced by our experiments (see Methods). Dispersion was highly correlated between species for single-copy (ρ=0.56) and multi-copy genes (ρ=0.42; Fig. 4C), suggesting that the determinants of expression variability are broadly conserved between *C. acutus* and *C. propinquus*. In both species, CSGs had much lower dispersion than non-DE and species-specific DE genes (LM, p<1 × 10^−4^; Fig. 5A). Among single-copy genes, CSGs had more similar dispersion estimates between species compared to non-DE and species-specific DE genes (LM, p<0.01; 5B); among multi-copy orthogroups, CSGs and species-specific DE genes both had more similar dispersion estimates between species than non-DE genes (LM, p<1 × 10^−4^). Thus, expression variability of CSGs was more conserved between *C. acutus* and *C. propinquus* than most other genes. In contrast, CHGs had higher dispersion than non-DE genes within species (LM, p<1 × 10^−4^; Table 3; Fig. S9) and no difference in dispersion between species.

**Figure 5:**
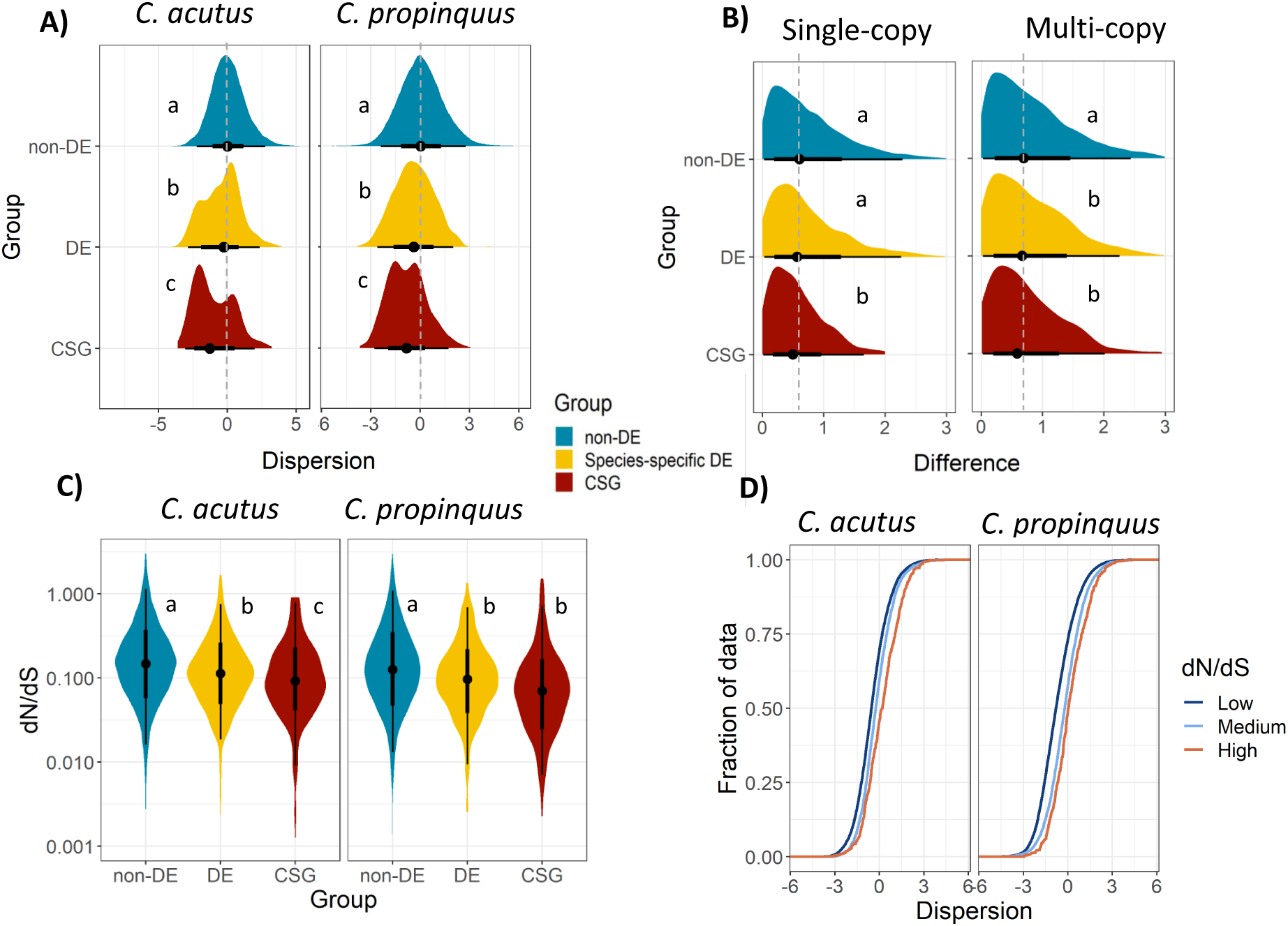
Starvation-response genes have less variable expression and lower rates of sequence evolution than other genes. (A) Distribution of dispersion estimates for *C. acutus* (left) and *C. propinquus* (right), calculated from field samples. Letters indicate significant differences between groups in a weighted linear regression (LM, p<0.05), and dashed lines indicate non-differentially expressed group median. For visualization, these are distributions of unweighted values. (B) Difference in dispersion estimates between species, for single-copy (left) and multi-copy (right) genes. For multi-copy genes, differences were calculated between average values of each gene family weighted by the inverse squared standard error of each gene’s dispersion estimate. (C) Branch-wise dN/dS estimates in *C. acutus* (left) and *C. propinquus* (right) calculated using gene trees inferred at the N0 node. Letters indicate significant differences between groups in a weighted logistic regression accounting for expression level (GLM, p<0.05). Note the log scale. (D) Empirical cumulative distribution function of dispersion estimates, calculated separately for genes with low (ω <0.2), medium (0.2 < ω < 0.8), and high (ω > 0.8) dN/dS estimates in the N0 dataset.

**Table 3:** Summary of expression and sequence characteristics of starvation-response genes (SGs) and handling-response genes (HGs) in copepods.

|  |  | non-DE | Species-specific<br>SGs | CSGs | Species-specific<br>HGs | CHGs |
| --- | --- | --- | --- | --- | --- | --- |
| Expression<br>level | *C. acutus* | 3.08<sup>a</sup> | 5.02<sup>b</sup> | 5.83<sup>c</sup> | 4.78<sup>d</sup> | 3.74<sup>e</sup> |
|  | *C. propinquus* | 3.47<sup>a</sup> | 5.63<sup>b</sup> | 4.93<sup>c</sup> | 5.58<sup>b</sup> | 4.06<sup>d</sup> |
| Expression<br>dispersion | *C. acutus* | -1.51<sup>a</sup> | -2.05<sup>b</sup> | -2.20<sup>c</sup> | -1.26<sup>d</sup> | -0.548<sup>e</sup> |
|  | *C. propinquus* | -1.32<sup>a</sup> | -1.52<sup>b</sup> | -1.94<sup>c</sup> | -1.22<sup>d</sup> | -0.766<sup>e</sup> |
| dN/dS | *C. acutus* | 0.233<sup>a</sup> | 0.191<sup>b</sup> | 0.154<sup>c</sup> | 0.206<sup>b</sup> | 0.240<sup>a</sup> |
|  | *C. propinquus* | 0.203<sup>a</sup> | 0.156<sup>b</sup> | 0.145<sup>b</sup> | 0.159<sup>b</sup> | 0.206<sup>a</sup> |
| Dispersion<br>divergence | Single-copy | 0.642<sup>a</sup> | 0.600<sup>a</sup> | 0.528<sup>b</sup> | 0.629<sup>a</sup> | 0.651<sup>a</sup> |
|  | Multi-copy | 0.944<sup>a</sup> | 0.627<sup>b</sup> | 0.540<sup>bc</sup> | 0.709<sup>c</sup> | 0.807<sup>a</sup> |

Letters indicate significant differences (GLM, $p < 0.05$ ; see Methods for statistical models) *within each row*. For expression level, values are mean TPM10k. For expression divergence, values are estimated marginal means of dispersion; negative values have lower variance than average after regressing out expression level. For dN/dS, values are estimated marginal means of dN/dS after accounting for expression level. For dispersion divergence, values are estimated marginal means of the absolute difference between dispersion across species. CSG: conserved starvation-response genes; CHGs: conserved handling-response genes; DE: differentially expressed.

### 3.5 Phylotranscriptomics support strong purifying selection on starvation-response genes

We compared branch-level dN/dS estimates using two datasets: 1) orthogroups inferred at the root of the species tree (N0), and 2) orthogroups inferred at the root of Calanoida (N1; see Fig. 1A). In both species and both datasets, starvation-response genes had lower dN/dS than other genes after accounting for expression level; in *C. acutus*, CSGs also had lower dN/dS than species-specific DE genes (GLM, p<0.01; Fig. 5C, Table 3). Thus, conserved and species-specific starvation-response genes appear to be under strong purifying selection at the sequence level. Species-specific handling-response genes had lower dN/dS than non-DE genes, but CHGs did not (Fig. S9). Across all genes, genes with lower dN/dS tended to have lower dispersion (ρ = 0.30; Fig. 5D).

We noticed that there was a subset of CSGs with particularly small LFC (<0.5), low dispersion, high expression, and low dN/dS. Even after excluding genes with abs(LFC) <0.5 from the analysis, CSGs still had lower dN/dS and more similar dispersion between species than other genes (at least in multi-copy orthogroups), although they no longer had lower dispersion within species. There was no such subset of small-LFC genes for the handling response.

To identify genes that may have experienced selection shifts in the OSC, we fit branch models of sequence evolution using codeml. In the N0 dataset, most (75%) gene families were better fit by a two-ratio model with separate dN/dS for OSC and background lineages (LRT, p<0.01), compared to a one-ratio model and a two-ratio model with OSC dN/dS set to 1 (neutral evolution). Strikingly, nearly all two-ratio genes (6352/6460, 98%) had lower dN/dS in the OSC, with a median reduction of 37% (Fig. 1B). In the N1 dataset, most (59%) genes were better fit by the two-ratio model, and the vast majority of these (6669/6852 97%) again had lower dN/dS in the OSC, with a median reduction of 34% (Fig. S10). This strongly suggests that purifying (negative) selection is pervasively stronger in the OSC compared to other copepods. For genes best fit by the two-ratio model, 2379 (38%) were better fit by as three-ratio model in which TAG-storing species (*C. propinquus* and *Eucalanus bungii*) were allowed a third dN/dS ratio (LRT, p<0.01). A total of 1857 (78%) of these genes had a higher dN/dS in TAG species than WE species. Across N0 genes, the median dN/dS was 0.252 in background lineages, 0.165 in WE lineages, and 0.179 in TAG lineages (Fig. 1B).

Against this backdrop of intensified selection, we found that gene families containing at least one CSG (n=232) had experienced a greater intensification of selection compared to families containing only species-specific DE (n=738) or non-DE (n=7643) genes (Fig. 1C; Fig. S11). This was true for both WE and TAG lineages, and the difference between CSGs and non-DE genes was slightly greater in WE compared to TAG species (GLM, p<0.05). The N1 dataset showed the same pattern, except with no significant difference between TAG and WE (Fig. S11). There was no such pattern for handling-response genes (Fig. S12).

We considered genes with increased dN/dS in the OSC as candidate targets of positive selection, and applied more sensitive tests in Hyphy. Using BUSTED-PH, just 18 genes (3/108 N0 in and 15/183 in N1) had evidence of positive selection specifically in the OSC (p<0.01; Table S3); none of these were DE. Using RELAX, we found that the vast majority of genes with elevated foreground dN/dS (102/108 for N0; 155/183 for N1) had experienced relaxed selection in the foreground (p<0.01). Therefore, elevated dN/dS of most of these genes can be explained by reduced intensity of purifying selection rather than invoking positive selection.

We manually annotated and investigated two key families of enzymes involved in lipid synthesis: fatty acid elongases (ELOVs) and desaturases (FADs). In our data, downregulated CSGs included one ELOV and one FAD orthogroup. Both species also downregulated additional (non-orthologous) FADs and *C. acutus* downregulated eight additional ELOVs (Fig. 6; Supplemental Information). We divided ELOVs into two ancestral paralogs for this analysis (see Methods). ELOV3/6 showed evidence of positive selection specifically in OSC lineages (BUSTED-PH, p<0.003), and relaxed selection in the OSC (RELAX, k=0.74, p<0.001); ELOV1/4/7 showed evidence of positive selection in both the foreground and background, and relaxed selection in the OSC (RELAX, k=0.85, p<0.001). FADs also had strong evidence of positive selection specifically in OSC species (BUSTED-PH, p<1× 10^−11^), and strongly relaxed selection in the OSC (RELAX, k=0.60, p<0.001). ELOVs and FADs appear to have expanded in the OSC, with higher median copy numbers in OSC taxa (medians of 12 ELOVs and 4 FADs in OSC compared to 6 ELOVs and 2 FADs in other species; Table S4) and gene tree topologies reflecting multiple duplications within this clade (Fig. 6).

**Figure 6:**
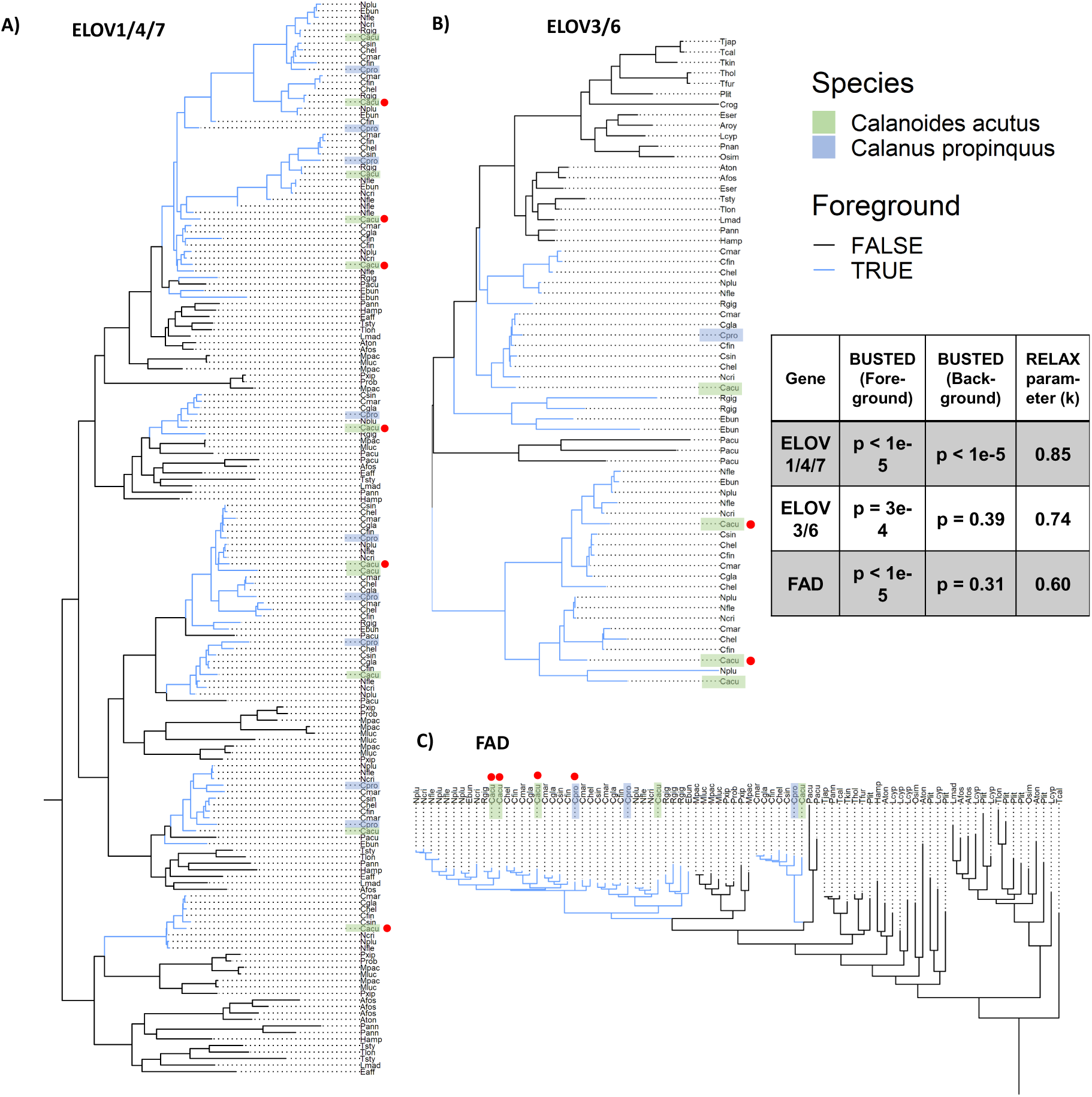
Gene trees of key copepod lipid enzymes. A) Fatty acid elongase ELOV1/4/7; B) ELOV3/6; C) Fatty acid desaturase (FAD); for clarity, we have zoomed in on a portion of the trees and full trees are available in the Supporting Information. Blue lines indicate species in the oil sac clade/foreground; *Calanoides acutus* and *Calanus propinquus* sequences are highlighted. Red dots indicate genes that were DE (downregulated) in starved animals. These trees were constructed from full-length transcripts (see Methods); partial-length transcripts not shown in these trees were still included in the expression analyses, and some partial-length transcripts were also DE (see text). Table indicates results of selection analyses (BUSTED-PH and RELAX). A significant BUSTED result indicates evidence for positive selection (at at least one site and branch); a value of k<1 indicates relaxation of selection in the foreground (all RELAX results were significant at p<0.001).

## 4 Discussion

The threat of starvation is particularly extreme for consumers in polar ecosystems due to strong seasonality and high patchiness in food distributions during the growing season (Smetacek and Nicol, 2005; Trudnowska et al., 2016). As such, high-latitude copepods often have higher starvation tolerance than tropical species, enabled by the acquisition of large energy reserves and metabolic strategies such as fall/winter diapause. Here, we studied the short-term (5-9 d) starvation responses of two species of Southern Ocean copepods (*Calanoides acutus* and *Calanus propinquus*) which, due to their large lipid stores and high biomass, form a critical component of Antarctic food webs. Using an equivalence test approach, we defined a conserved starvation response consisting of homologs with similar responses to starvation between species. Importantly, this allowed us to propagate uncertainty in fold-change estimates and control the false discovery rate at the level of orthologs. The commonly-employed “Venn diagram” approach of selecting the overlap between lists of DE genes is too conservative and would include many false negatives (consider a gene with p=0.049 in one species and p=0.051 in the other; Gelman and Stern, 2006).

The conserved starvation response included changes to genes that mediate RNA metabolism, protein translation and degradation, and DNA metabolism and repair, as well as vitellogenins and some enzymes and transcription factors with key roles in lipid metabolism. In general, genes related to protein metabolism responded similarly to starvation across species while lipid metabolism genes differed; this probably reflects the different lipid storage strategies of *C. acutus* (WE storage) and *C. propinquus* (TAG storage). Starved animals in both species also upregulated genes related to the cell cycle and stress response, and downregulated genes related to immune response, oxidoreductase activity, and carbohydrate metabolism (though these genes were not necessarily homologous). Together, these gene expression changes are consistent with reduced lipid synthesis, altered cellular metabolism, changes to mitochondrial and energy metabolism, and modulation of developmental and reproductive processes.

Although OSC species can survive weeks without food (Helland et al., 2003; Lee et al., 2006), our experiment was long enough to observe some physiological changes in lipid content and enzyme activity. Starved *C. acutus* had less lipid than when fed, although our sample size was limited. There was no difference in lipid content of starved *C. propinquus* compared to fed. However, all *C. propinquus* in our study were lipid-poor, with more membrane phospholipids than storage TAG. This is not unusual, as *C. propinquus* in other parts of the Southern Ocean also have low lipid content in December-January, probably because females have allocated their resources towards reproduction instead of lipid storage (Hagen and Schnack-Schiel, 1996; Pasternak et al., 2009; Graeve et al., 2020). Both species exhibited reduced citrate synthase activity following starvation and concordant downregulation of genes involved in aerobic respiration, indicating reduced aerobic metabolic potential. This is similar to other calanoid copepods, where citrate synthase activity correlates with food availability on 2-3 day time scales (Clarke and Walsh, 1993).

Our results suggest that some of the same regulatory pathways likely mediate starvation-induced changes to protein and lipid metabolism in copepods as in insects. Protein synthesis is one of the most energy-intensive cellular processes (Buttgereit and Brand, 1995), and therefore an overall decrease in protein synthesis and ribosome production is a critical step in tolerating food deprivation. In *Drosophila*, starvation reduces insulin-like signalling, modulating transcription factors such as FoxO and CREB and leading to the upregulation of lipases that hydrolyze triglycerides (DiAngelo and Birnbaum, 2009; Lehmann, 2018). This supplies energy to the animal. Insulin signalling also activates the TOR pathway, and changes in insulin/TOR signalling are associated with the inhibition of protein synthesis during starvation (Terashima and Bownes, 2005; Hietakangas and Cohen, 2009). Consistent with this, the conserved starvation response in *C. acutus* and *C. propinquus* was characterized by genes that mediate ribosome biogenesis, translation, and protein catabolism. Both species upregulated a homolog of TOR, reflecting its central role in nutrient sensing. Each species also modulated additional transcription factors and genes related to insulin signalling, particularly *C. propinquus*. Changes in insulin signalling may be more apparent in *C. propinquus* because it stores TAG, whereas the wax ester storage of *C. acutus* may be regulated by other genes.

Several factors resulting from inherent challenges of ship-board experiments should be kept in mind when interpreting our results. We noted substantial gene expression changes between (fed) experimental samples and field samples, indicating a stress response to captivity. Consistent with this, *C. acutus* (but not *C. propinquus*) exhibited some mortality throughout the experiment in both fed and starved treatments. Additionally, some starved animals had visible food in their guts (Fig. S4). Algal growth during the experiments is unlikely because they were kept in the dark. However, due to logistical constraints, we could not always remove some small algal cells (<3 microns) from the starved treatments. Nonetheless, changes in physiological condition, gene expression, and the correlation of our experimental response with field chlorophyll measurements reported in Berger et al. (2023) indicate that our starvation treatment captured a *bona fide* response to food deprivation.

### 4.1 Conserved starvation-response genes (CSGs) are under purifying and stabilizing selection

Starvation-response genes are under strong constraint at both the expression and sequence levels, suggesting that they are critical for organismal fitness. Starvation-response gene sequences are under strong purifying selection even after accounting for expression level, based on branch-level dN/dS estimates in our focal species. Clade-level analyses also found evidence for low dN/dS in CSGs across the copepod phylogeny. Starvation-response genes had less variable expression than other genes in our paired field study (Berger et al., 2023) while having similar dispersion across species, strongly suggesting the action of stabilizing selection on expression level (Price et al., 2022). While purifying selection on sequence and stabilizing selection on expression need not go hand in hand, we found that genes under strong coding constraint tend to have less expression variability among individuals (Fig. 5D). This is consistent with similar results in primates (Fair et al., 2020), suggesting that coding sequence conservation and expression variability are complementary indicators of selective constraints on gene products. The starvation response was more similar between species than the handling response in terms of gene expression changes, expression variability, and sequence conservation (Table 3), indicating that starvation-response genes are under particularly strong sequence and expression constraint even compared to other stress-response genes.

The conserved starvation response included a set of upregulated genes with small (<0.5) LFC in both species. Importantly, there was no comparable set of handling-response genes with small LFC, meaning their presence in the starvation dataset is unlikely to be an artifact of the DE analysis. RNA-seq analyses commonly ignore genes below some fold-change cutoff, with the justification that genes with large fold-change are more biologically relevant. However, choice of a cutoff is arbitrary and biologically relevant genes can exhibit small fold-changes for many reasons (St. Laurent et al., 2013). For instance, bulk RNA-seq measurements dilute tissue-specific responses, such as may occur in the gut or oil sac during starvation. Qualitatively, small-LFC genes included many genes with high expression and fundamental roles in cellular processes, such as ribosome and proteasome components, helicases and polymerases, and translation initiation factors. Since these genes are highly pleiotropic and essential but may nonetheless be modulated by energy restriction, their expression may be precisely regulated during starvation, resulting in the observed small-but-precise fold-changes. These small-LFC genes contribute to the strong selection pressure observed on starvation-response genes, but we note that our inferences of reduced dispersion between species and low dN/dS (at least in *C. acutus*) were robust to their exclusion.

### 4.2 Implications for the evolution of the oil-sac clade (OSC)

Although it is not the main focus of our paper, we are the first to construct an order-level copepod phylogeny from genome-scale data. Due to biased taxon sampling, low completeness of some assemblies, and the vast evolutionary distances involved, we took several steps to minimize systematic bias and potential sources of long-branch attraction (see Methods). We recovered major clades of copepods with strong support, including known superfamily-level relationships within Calanoida (Blanco-Bercial et al., 2011). Unlike some previous studies, we place *Calanoides* sister to *Neocalanus*+*Calanus* rather than recovering *Calanoides*+*Neocalanus* (Bucklin et al., 2003); this relationship might be clarified by sampling additional species. Several parts of the phylogeny are still difficult to resolve (namely the placement of *Platychelipus* and relationships among Temoridae, Pontellidae, and Acartiidae), although this does not affect our other results. Sequencing additional high-quality transcriptomes, especially outside Calanoida, is needed to better understand copepod phylogenetics.

We focused our evolutionary analyses on three families characterized by large body size, large oil sacs, and diapause (Calanidae+Eucalanidae+Rhincalanidae). These taxa are united by their ecology, physical characteristics, and general metabolic strategy, and they play a unique role in temperate and polar ecosystems. For ease of reference, we termed this group the “oil sac clade” (OSC); however, it is important to note that oil sacs and diapause/dormancy are present to various extents in other, distantly related copepod taxa (e.g. *Metridia spp.*, Hays et al., 2001). We are not aware of any study explicitly examining the distribution or homology of oil sac structures among copepods, and evolutionary analyses of these traits might be an interesting area for future study.

We leveraged transcriptomic resources available for copepods to infer selection pressure on genes in the OSC compared to other lineages. Surprisingly, transcriptome-wide dN/dS was substantially lower in the OSC compared to other copepods, and even compared to other calanoids. One plausible explanation is higher effective population sizes of OSC taxa, which include some of the most abundant animal species on Earth (Humes, 1994). However, other calanoid copepods, such as *Oithona spp.*, are also extremely abundant, and estimates of effective population size in pelagic copepods are rare (but see Bucklin and Wiebe, 1998; Provan et al., 2008).

CSGs (defined in *C. acutus* and *C. propinquus*) appear to be under strong purifying selection across the copepod phylogeny, but the relative strength of selection on these genes has increased in OSC lineages, and particularly in WE-storing species (Fig. 2C; Fig. S11). In contrast, patterns of selection on handling-response genes did not differ from the rest of the transcriptome (Fig. S12). One explanation for this pattern is that CSGs mediate an ancestral starvation response present in other copepod lineages and that selection pressure on starvation tolerance itself has intensified in the OSC, possibly due to the regular exposure of these taxa to long periods of food deprivation. This is plausible given the enrichment of CSGs for functions known to mediate starvation tolerance in other arthropods. The difference between WE-storing and TAG-storing species is also consistent with the generalization that WE-storing species are more reliant on herbivory and long-term lipid storage rather than opportunistic year-round feeding. However, another possibility is that some CSGs acquired a role in the starvation response in the OSC and subsequently became subject to stronger selection.

Our initial analysis did not identify a signal of positive selection in the OSC associated with starvation-response genes or energy metabolism, but was limited by the use of codeml branch models to identify candidate genes—as branch-site tests were too computationally intensive to apply to the whole transcriptome—and by reliance on orthogroups defined by Orthofinder, which tend to over-split large gene families. However, we reasoned that the evolution of OSC taxa likely involved modification or redeployment of lipid metabolism pathways, so we manually annotated two key gene families involved in lipid metabolism, fatty acid elongases (ELOVs) and desaturases (FADs). We found evidence for positive (diversifying) selection within these families in OSC taxa.

ELOVs catalyze the synthesis of very long-chain fatty acids (Castro et al., 2016), while FADs introduce double bonds into long-chain fatty acids and are critical for the production of membrane phospholipids and storage lipids (Nakamura and Nara, 2004). In OSC species, both families of enzymes are upregulated during juvenile development and preparation for diapause (Tarrant et al., 2008, 2014; Lenz et al., 2014; Berger et al., 2021) and downregulated in low-food conditions (Roncalli et al., 2019, 2022), consistent with their role in lipid synthesis. ELOV and FAD genes were downregulated in starved *C. acutus* and *C. propinquus*, with different copies displaying either conserved or species-specific responses to starvation. We found evidence for positive selection acting on ELOVs and FADs within the OSC, along with relaxed selection suggesting reduced functional constraint. Our selection analyses, along with multiple apparent duplications of both enzymes in OSC lineages (Fig. 6), suggest a scenario in which these enzymes repeatedly duplicated in the OSC followed by a relaxation of selective constraints and possible sub- or neo-functionalization. Lineage-specific duplications have played a major role in the distributions of elongases and desaturases in a variety of taxa (Surm et al., 2015, 2018; Ishikawa et al., 2019), and copy number expansion of ELOVs has been suggested to contribute to the ability of some harpacticoid copepods to synthesize long-chain fatty acids (Boyen et al., 2020). Higher copy numbers might facilitate the production of OSC species’ enormous lipid stores, while diversifying selection implies that some duplicated gene copies have evolved new functions such as distinct substrate specificities, as has occurred with vertebrate elongases (Castro et al., 2016). Biochemical analyses could test this hypothesis and are an intriguing area for future work.

## 5 Data Availability

Raw RNA-seq data have been uploaded to the NCBI Sequence Read Archive (SRA) and transcriptomes have been uploaded to the Transcriptome Shotgun Assembly Database (TSA), with Bioprojects PRJNA757455 (Calanoides acutus) and PRJNA669816 (Calanus propqinuus). R code used for analysis is available at https://github.com/caberger1/Antarctic_copepod_scripts

## 6 Acknowledgments

We would like to thank Dr. Junya Hirai for generously providing sequencing data for *Eucalanus bungii* prior to public release, Dr. Leocadio Blanco-Bercial for advice and comments regarding the phylogenetic analysis, and Dr. Joe Ryan for discussion. We also thank the Captain, crew and technical staff of the ARSV *Lawrence M. Gould*; Joe Cope, Dr. Patricia Thibodeau, and other members of the Steinberg Lab for copepod sampling and other assistance at sea; Dr. Mark Baumgartner and Dr. Phil Alatalo for technical assistance; and Nancy Copley and Adrienne Jones for assistance in photo analysis. We thank Ms. Michelle Stowell for her assistance with lipid classes analyses at the Marine Lipid Ecology Lab at the Hatfield Marine Science Center. Funding for this project was provided by the National Science Foundation Office of Polar Programs (Grants OPP-1746087 to AMT, and OPP-1440435 and OPP-2026045 to DKS).

## 7 Author Contributions

DKS and AMT conceived and designed the research. AMT collected the copepods and performed the feeding experiments. LAC collected and analyzed the lipid data, and AMT collected the other physiological measurements. CAB analyzed the sequencing data and performed evolutionary analyses. CAB and AMT wrote the manuscript with input from DKS and LAC, and all authors read and approved the final version.

