## Supplemental text, figures, and tables for "Comparative analysis of the molecular starvation response of Southern Ocean copepods"

Cory A. Berger, Deborah K. Steinberg, Louise A. Copeman, Ann M. Tarrant

**Table of Contents:**

| **Supplemental Text** | Page 2 |
| --- | --- |
| **Supplemental Tables** | Page 6 |
| **Supplemental Figures** | Page 10 |

**File S2: full differential expression and gene ontology enrichment results, WGCNA module assignments, gene annotations, and orthogroup assignments.**

**File S3: WGCNA GO enrichment, *Calanoides acutus***

**File S4: WGCNA GO enrichment, *Calanus propinquus***

**File S5: Full phylogenetic tree, ELOV1/4/7**

**File S6: Full phylogenetic tree, ELOV3/6**

**File S7: Full phylogenetic tree, FAD**

**Supplemental Text**

***Citrate synthase activity***

Copepods were thawed on ice, blotted on a lint-free tissue (Kimwipe, Kimberly-Clark, Irving TX), and quickly weighed on a Cahn C-33 microbalance. Groups of 2-5 copepods were pooled into 300 µL of ice-cold homogenization buffer (25 mM Tris, pH 7.8, 1mM EDTA, 10% glycerol) in a 5-mL Potter-Elvehjem homogenizer. Copepod tissue was homogenized using a motorized PTFE pestle for two 30-second bursts with 30 seconds of ice cooling between bursts. Homogenates were centrifuged at 14,000g for 20 minutes at 4°C, and the supernatant was retained. Enzyme activity measurements were made with 20 µL homogenate per well in triplicate wells of a 96-well plate. Measurements were made at 26°C using a SpectraMax plate reader. The automix function was used prior to each set of measurements. CS activity was measured modifying the protocol of Hawkins et al. (2016). 170 µL of CS assay buffer (0.11% Triton X-100, 294 μM 5,5’-dithiobis-[2-nitrobenzoic acid] [DTNB], 588 μM acetyl-coenzyme A in CS homogenization buffer, made fresh daily) was added to the homogenate. After taking baseline absorbance measurements for 3 minutes at 405 nm, 10 µL of 10 mM oxaloacetate was added to each well, and the change in absorbance at 405 nm was recorded over 3 minutes.

CS activity was determined by comparing measurements of copepod homogenates with a standard curve derived from a dilution series of a pure enzyme standard (CS from porcine heart, Sigma-Aldrich). Enzyme activity was normalized to wet mass.

***Lipid quantification***

Samples were processed over ice with a dissecting microscope to enumerate the number of individual zooplankton per sample. Individuals were then placed on a Kimwipe to remove excess moisture and were weighted on a microbalance in order to determine the wet mass of the lipid sample (+/- 0.0001 mg). Tissues were homogenized in chloroform and methanol and total lipids were extracted according to Parrish (1987) using a modified Folch procedure (Folch et al. 1956).

Total lipids were determined using thin layer chromatography with flame ionization detection (TLC/FID) with a MARK VI Iatroscan (Iatron Laboratories, Tokyo, Japan) as described by Lu et al. (2008) and Copeman et al. (2017). Extracts were spotted on duplicate silica-gel-coated Chromarods, and a three-stage development system was used to separate wax esters, triacylglycerols, free fatty acids, sterols and polar lipids. Polar lipid is mostly comprised of phospholipids with minor amounts of other acetone mobile polar lipids. The first rod development was in a chloroform: methanol: water solution (5:4:1 by volume) until the leading edge of the solvent phase reached 1 cm above the spotting origin. The rods were then developed in hexane: diethyl ether: formic acid solution (99:1:0.05) for 38 min, and finally rods were developed in a hexane: diethyl ether: formic acid solution (80:20:0.1) for 38 min. After each solvent development, rods were dried (5 min) and conditioned (5 min) in a constant humidity chamber (~32%) that was saturated with aqueous CaCl2. Following the last development, rods were scanned using Peak Simple software (ver. 3.67, SRI Inc.) and the signal detected in millivolts was quantified with calibration curves using the following standards from Sigma (St Louis, MO, USA): palmitic acid (free fatty acids), cholesterol (sterols), L-alpha-phosphatidylcholine (polar lipids). Specialized standards were purified by column chromatography to use for wax esters (*Calanus finmarchicus* oil) and triacylglycerols (*Boreogadus saida* liver) using methods from Ohman (1997). Calibrated relationships between lipid class areas and standard lipid amounts (µg) had correlations with an r2 >0.98 for all classes.

Following total lipid and lipid class analyses of zooplankton tissues, samples were processed for fatty acid and alcohol analyses. All samples and total lipid extracts were derivatized into their fatty acid methyl esters (FAMEs) and free fatty alcohols using sulphuric acid-catalyzed transesteriﬁcation (Budge et al. 2006). Resulting FAMEs and alcohols were simultaneously analyzed on a DB-FFAP GC column (Agilent Technologies, Inc., U.S.A.). The column was 30 m in length, with an internal diameter of 0.25 mm and a film thickness of 0.25. The column temperature began at 150 °C and held this temperature for 0.5 min. Temperature was then increased to 250 °C using a ramp of 4 °C min -1 and this final temperature was held for 8 min. The total run time was 30 minutes. The carrier gas was helium, flowing at a rate of 1 ml min-1. The injector temperature was set at 250 °C and the detector temperature was constant at 250 °C (Kattner & Fricke 1986, Graeve et al. 2005). Peaks were identified using retention times based upon standards purchased from Supelco (37 component FAME, BAME, PUFA 1, PUFA 3). Column function was checked by comparing chromatographic peak areas to empirical response areas using a quantitative FA mixed standard, GLC 487 (NuCheck Prep). Chromatograms were integrated using Chem Station (version A.01.02, Agilent).

***Phylotranscriptomics***

Table S1 contains information about reads used for each transcriptome assembly. For species where only the raw reads were available or the existing assembly was of low quality, we assembled a new transcriptome from raw reads using a common trimming and assembly pipeline, which was identical to the procedure used to assemble *C. acutus* (kmer correction with RCorrector, quality trimming with TrimGalore with a Phred score cutoff of 5, rRNA removal with SortMeRNA, and assembly with Trinity). We ran OrthoFinder v2.5.2 (Emms & Kelly, 2019) on the dataset of 41 transcriptomes to infer gene families.

OrthoFinder identified 1355 genes that were single-copy in at least 22\% (9/41) of species. Within each OrthoGroup, we aligned genes using MAFFT L-INS-I v7.475 (Katoh et al.,

2009) and built a maximum-likelihood (ML) tree using IQ-TREE v2.1.4 (Nguyen et al.,

2015) with the best-fit model according to ModelFinder (Kalyaanamoorthy et al., 2017). We used TreeShrink (Mai & Mirarab, 2018) with alpha=0.05 to remove unrealistically long branches (which may represent errors in alignment or orthology assignment), re-aligned the remaining sequences, used Trimal v1.2 (Capella-Gutiérrez et al., 2009) to remove sites with less than 20% occupancy, and dropped genes less than 50 amino acids long after filtering, resulting in a dataset of 1001 genes. We used phykit (Steenwyk et al., 2021) to calculate the LB score of each gene tree (Struck et al, 2014), which is a measure of branch-length heterogeneity, and retained the 50% of genes with the lowest LB scores. This resulted in a set of 500 genes, containing 88,379 sites, that should be less susceptible to long-branch attraction. We applied an edge-proportional partition analysis (Chernomor et al., 2016) to this dataset with IQ-TREE, with the optimal partitioning scheme inferred by PartitionFinder (Lanfear et al., 2021). We used the partition tree as the guide tree for a posterior mean site frequency model (PMSF; Wang et al., 2018), Q.pfam+C20+F+R5. We retained the best tree from 5 independent runs, and assessed branch support with ultra-fast bootstraps (Hoang et al., 2018) and SH-aLRT (Guindon et al., 2010) with 1000 replicates each. The partition and PMSF approaches recovered identical topologies. We rooted the tree between Calanoida (Gymnoplea) and the other orders (Podoplea), which is well-supported by previous morphological and molecular studies (Huys & Boxshall, 1991; Huys et al., 2007; Eyun, 2017).

**Supplemental Tables**

**Table S1.** Details of copepod transcriptomes used in this study (see file “Copepod_sources.xls’”).

**Table S2.** Accession numbers of representative metazoan fatty acid elongases (ELOV) and desaturases (FAD) used for homology search.

| Species | GenBank Accession | Gene |
| --- | --- | --- |
| *Homo sapiens* | AYN74362.1 | ELOV |
| *Homo sapiens* | Q9NXB9.2 | ELOV |
| *Homo sapiens* | Q9HB03.2 | ELOV |
| *Homo sapiens* | NP_073563.1 | ELOV |
| *Homo sapiens* | Q9NYP7.1 | ELOV |
| *Homo sapiens* | NP_001124193.1 | ELOV |
| *Homo sapiens* | A1L3X0.1 | ELOV |
| *Danio rerio* | NP_001315523.1 | ELOV |
| *Danio rerio* | XP_005162628.1 | ELOV |
| *Danio rerio* | AAI52204.1 | ELOV |
| *Danio rerio* | NP_956747.1 | ELOV |
| *Danio rerio* | NP_955826.1 | ELOV |
| *Danio rerio* | NP_956169.1 | ELOV |
| *Danio rerio* | NP_001070061.1 | ELOV |
| *Danio rerio* | NP_001019609.2 | ELOV |
| *Gallus gallus* | XP_015146466.1 | ELOV |
| *Gallus gallus* | NP_001184237.1 | ELOV |
| *Gallus gallus* | NP_001305339.1 | ELOV |
| *Gallus gallus* | NP_001184238.1 | ELOV |
| *Gallus gallus* | XP_015140326.1 | ELOV |
| *Gallus gallus* | NP_001026710.1 | ELOV |
| *Gallus gallus* | NP_001184239.1 | ELOV |
| *Drosophila melanogaster* | NP_729666.2 | ELOV |
| *Drosophila melanogaster* | NP_001097580.1 | ELOV |
| *Drosophila melanogaster* | AAF54172.2 | ELOV |
| *Drosophila melanogaster* | Q9VH58.1 | ELOV |
| *Drosophila melanogaster* | NP_651062.3 | ELOV |
| *Drosophila melanogaster* | Q9VV87 | ELOV |
| *Drosophila melanogaster* | NP_001246905.2 | ELOV |
| *Scylla olivacea* | QNV46824.1 | ELOV |
| *Scylla olivacea* | AWM30548.1 | ELOV |
| *Scylla olivacea* | QBX90561.1 | ELOV |
| *Penaeus vannamei* | ROT84478.1 | ELOV |
| *Penaeus vannamei* | ANK58182.1 | ELOV |
| *Penaeus vannamei* | AKJ77890.1 | ELOV |
| *Daphnia magna* | KZS12713.1 | ELOV |
| *Daphnia magna* | KZS11353.1 | ELOV |
| *Daphnia magna* | XP_032796449.1 | ELOV |
| *Daphnia magna* | XP_032778990.1 | ELOV |
| *Homo sapiens* | O60427 | FAD |
| *Homo sapiens* | O95864 | FAD |
| *Homo sapiens* | Q9Y5Q0 | FAD |
| *Danio rerio* | NP_571720.2 | FAD |
| *Gallus gallus* | XP_421052.4 | FAD |
| *Gallus gallus* | XP_426408.2 | FAD |
| *Gallus gallus* | NP_001153900.1 | FAD |
| *Drosophila melanogaster* | NP_652731.1 | FAD |
| *Drosophila melanogaster* | NP_650201.1 | FAD |
| *Amphibalanus amphritrite* | KAF0313635.1 | FAD |
| *Daphnia magna* | XP_032783737.1 | FAD |
| *Daphnia magna* | KZS12481.1 | FAD |
| *Armadillidium vulgare* | RXG58861.1 | FAD |
| *Penaeus vannamei* | XP_027230509.1 | FAD |
| *Scylla olivacea* | QKG32713.1 | FAD |
| *Sepia officinalis* | AKE92955.1 | FAD |
| *Nematostella vectensis* | XP_001640617.1 | FAD |
| *Actinia tenebrosa* | XP_031565068.1 | FAD |
| *Acropora millepora* | XP_029179496.1 | FAD |
| *Acropora millepora* | XP_029179499.1 | FAD |

**Table S3.** Orthogroups with evidence of positive selection in the OSC (BUSTED-PH*)

| Orthogroup | Annotation |
| --- | --- |
| N0.HOG0026672 | unannotated/shisa-like |
| N0.HOG0006594 | unannotated |
| N0.HOG0036752 | Methyl-CpG-binding domain protein |
| N1.HOG0017687 | BAZ2B |
| N1.HOG0030463 | DNA helicase (RECQ4) |
| N1.HOG0044818 | BRC1 (Broad-complex protein) |
| N1.HOG0024179 | unannotated |
| N1.HOG0029442 | peroxiredoxin |
| N1.HOG0026276 | tyrosine protein kinase |
| N1.HOG0043419 | ER membrane protein? |
| N1.HOG0021931 | CD151 antigen / tetraspanin |
| N1.HOG0019575 | Potassium channel subfamily K |
| N1.HOG0041377 | unannotated |
| N1.HOG0010900 | Complement C1q-like |
| N1.HOG0039554 | unannotated |
| N1.HOG0040243 | unannotated |
| N1.HOG0018430 | unannotated |
| N1.HOG0039149 | Transcription factor hamlet |

*(p*>*0.01 in foreground, p*>*0.05 in background, p*<*0.01 for difference between foreground and background). See Supplemental File 2 for full annotations of these and other genes.

**Table S4.** Copy numbers of ELOVs and FADs identified in copepod transcriptomes.

| Species | # ELOV1/4/7 | # ELOV3/6 | # FAD |
| --- | --- | --- | --- |
| *Acartia fossae* | 6 | 1 | 2 |
| *Acartia tonsa* | 2 | 1 | 10 |
| *Apocyclops royi* | 4 | 1 | 2 |
| *Calanoides acutus* | 13 | 3 | 5 |
| *Calanus finmarchicus* | 12 | 4 | 4 |
| *Calanus glacialis* | 7 | 2 | 4 |
| *Calanus helgolandicus* | 9 | 5 | 3 |
| *Calanus marshallae* | 11 | 4 | 5 |
| *Calanus propinquus* | 10 | 1 | 3 |
| *Caligus rogercresseyi* | 6 | 1 | 2 |
| *Calanus sinicus* | 7 | 2 | 3 |
| *Eucalanus bungii* | 7 | 3 | 2 |
| *Eucyclops serrulatus* | 8 | 1 | 2 |
| *Eurytemora affinis* | 5 | 1 | 0 |
| *Hemidiaptemus amplyodon* | 5 | 1 | 2 |
| *Labidocera madurae* | 4 | 1 | 1 |
| *Lepeophtheirus salmonis* | 7 | 0 | 1 |
| *Lernaea cyrpinacea* | 5 | 1 | 16 |
| *Metridia lucens* | 7 | 0 | 2 |
| *Metridia pacifica* | 9 | 0 | 3 |
| *Mytilicola intestinalis* | 0 | 0 | 0 |
| *Neocalanus cristatus* | 7 | 3 | 3 |
| *Neocalanus flemingerii* | 10 | 4 | 4 |
| *Neocalanus plumchrus* | 9 | 4 | 6 |
| *Oithona nana* | 2 | 0 | 0 |
| *Oithona similis* | 2 | 1 | 5 |
| *Paracyclopina nana* | 3 | 1 | 0 |
| *Platychelipus littoralis* | 8 | 1 | 31 |
| *Pleuromamma robusta* | 2 | 0 | 1 |
| *Pleuromamma xiphias* | 6 | 0 | 2 |
| *Pseudocalanus acuspes* | 6 | 3 | 2 |
| *Pseudodiaptemus annandalei* | 6 | 1 | 1 |
| *Rhincalanus gigas* | 6 | 3 | 4 |
| *Temora longicornis* | 3 | 1 | 3 |
| *Temora stylifera* | 6 | 1 | 0 |
| *Tigriopus californicus* | 6 | 1 | 6 |
| *Tigriopus japonicus* | 5 | 1 | 5 |
| *Tigriopus kingsejongensis* | 6 | 1 | 5 |
| *Tisbe furcata* | 2 | 1 | 3 |
| *Tisbe holothuriae* | 5 | 1 | 2 |
| *Tracheliastes polycolpus* | 5 | 0 | 1 |

**Supplemental Figures**

A)

B)
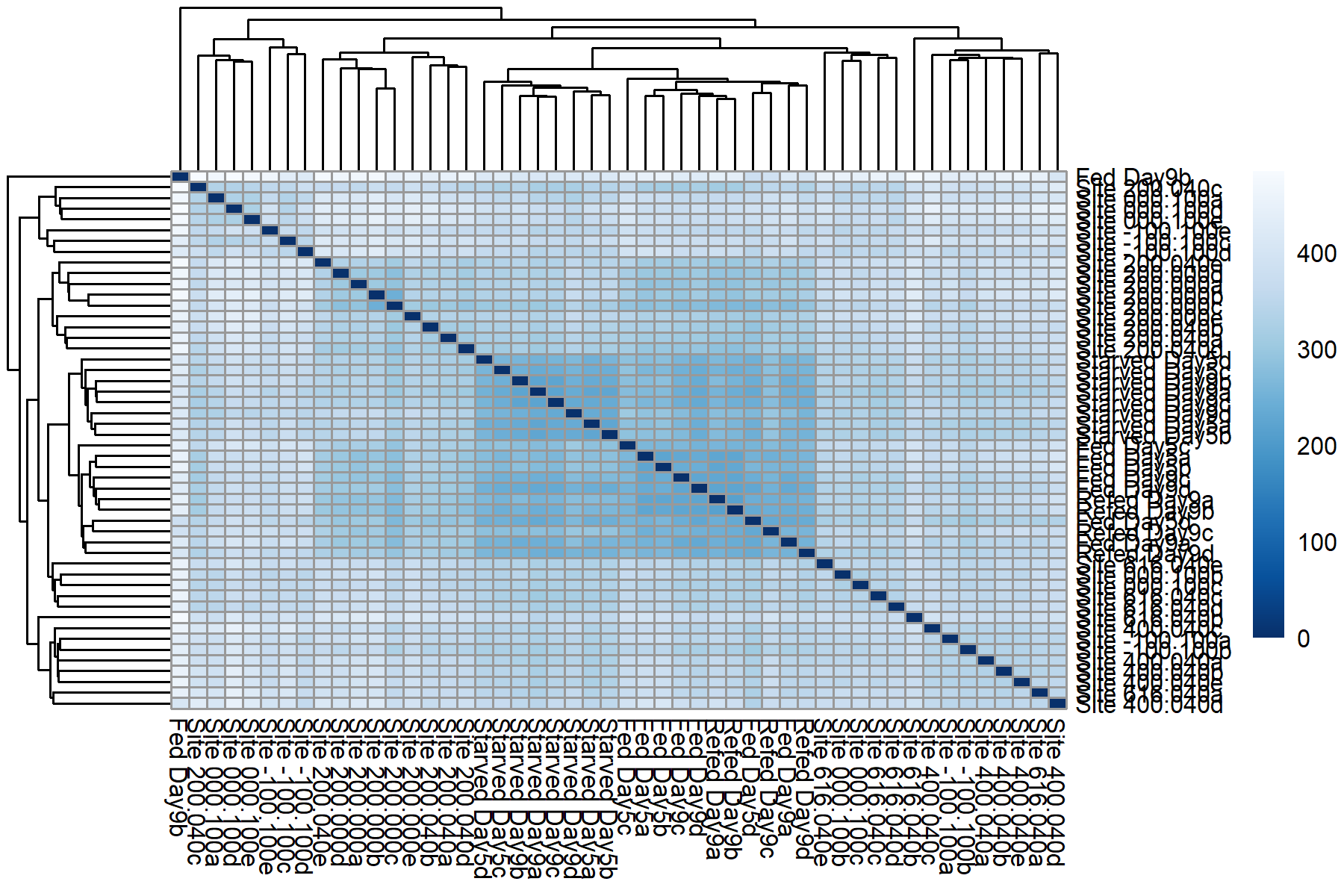

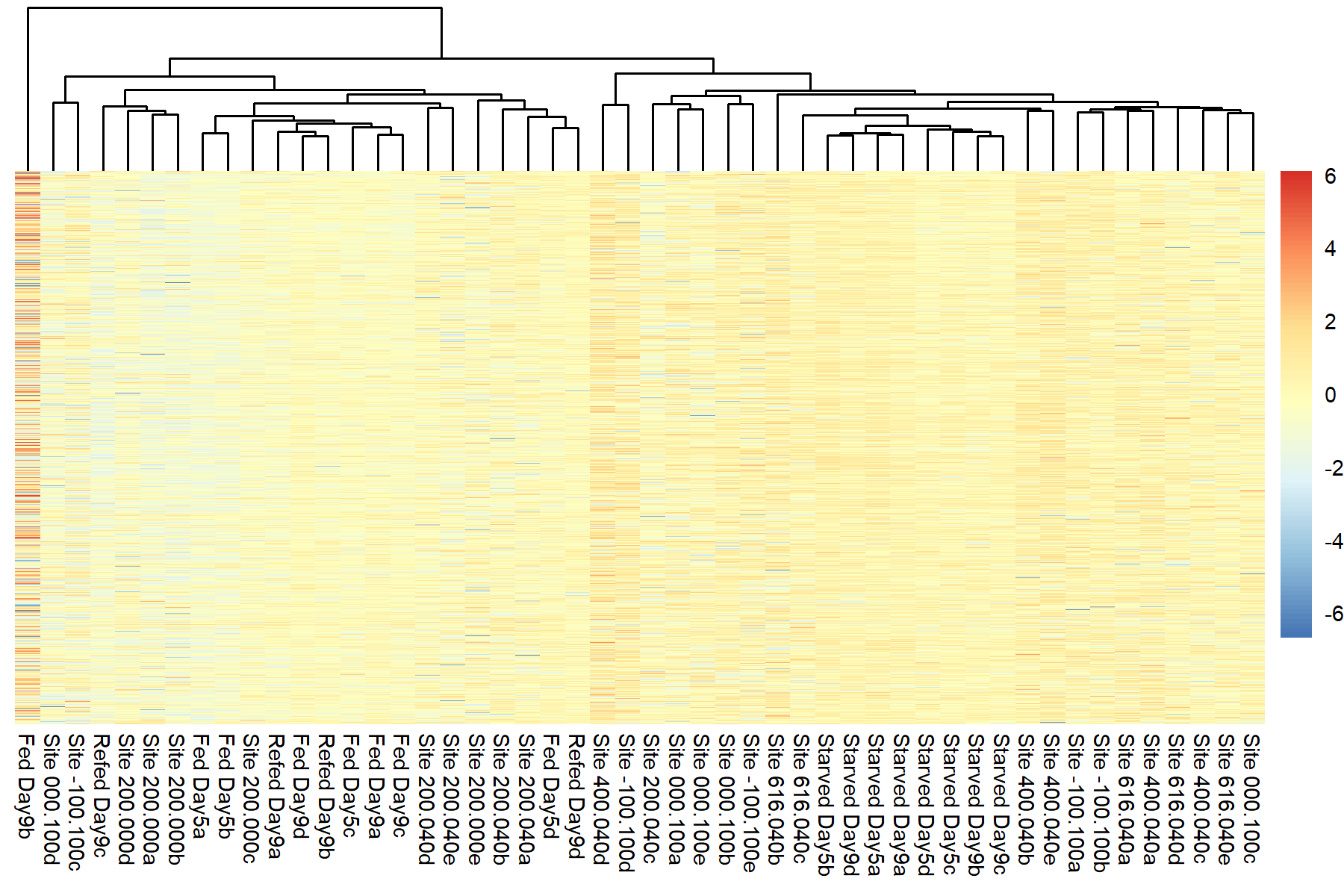

**Fig. S1.** Sample clustering based on sample distance matrix (A) and scaled and centered expression of the top 5000 most highly expressed genes (B) informed our decision to remove the outlier sample ‘Fed Day9b’ in *Calanoides. acutus*.

A)

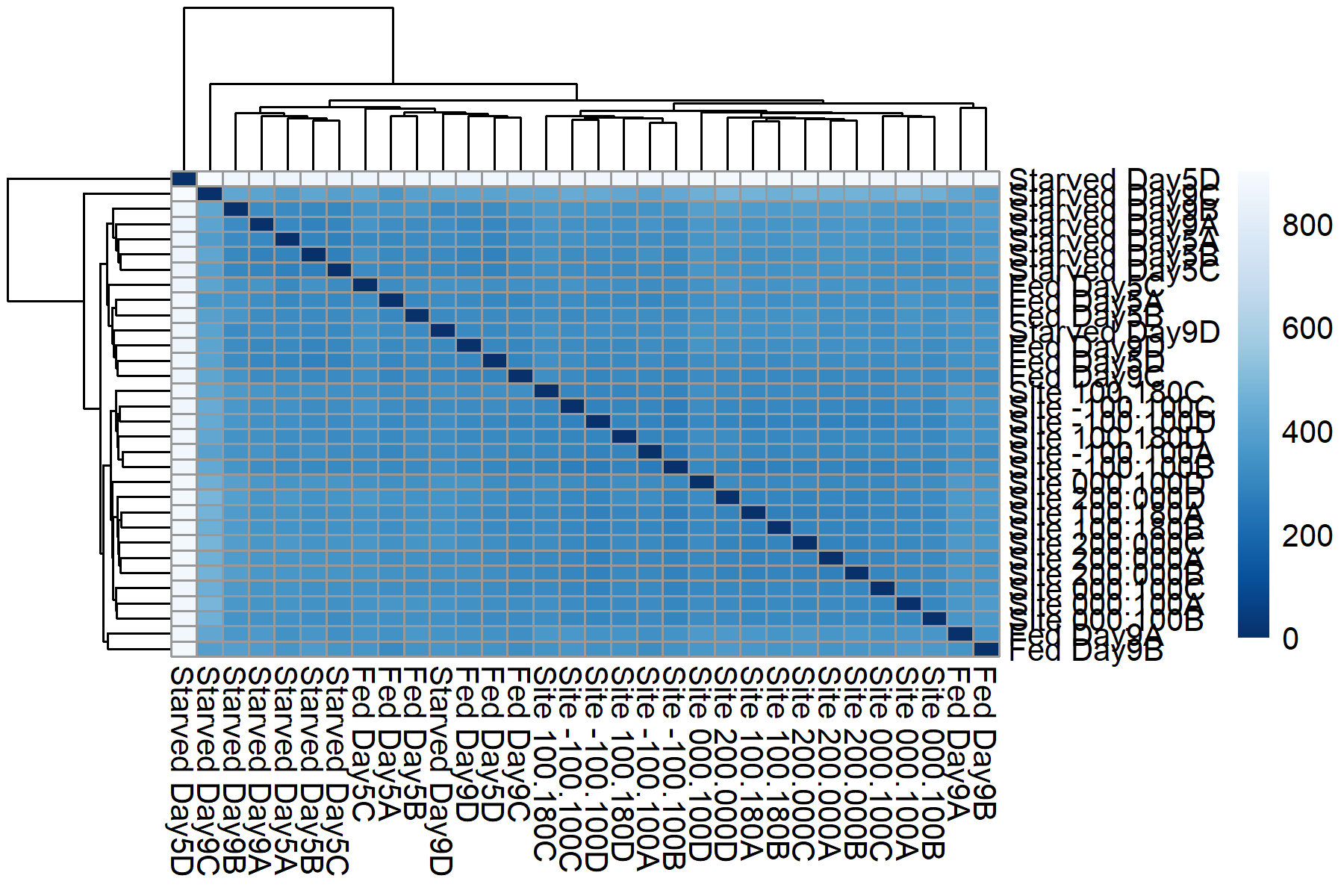

B)

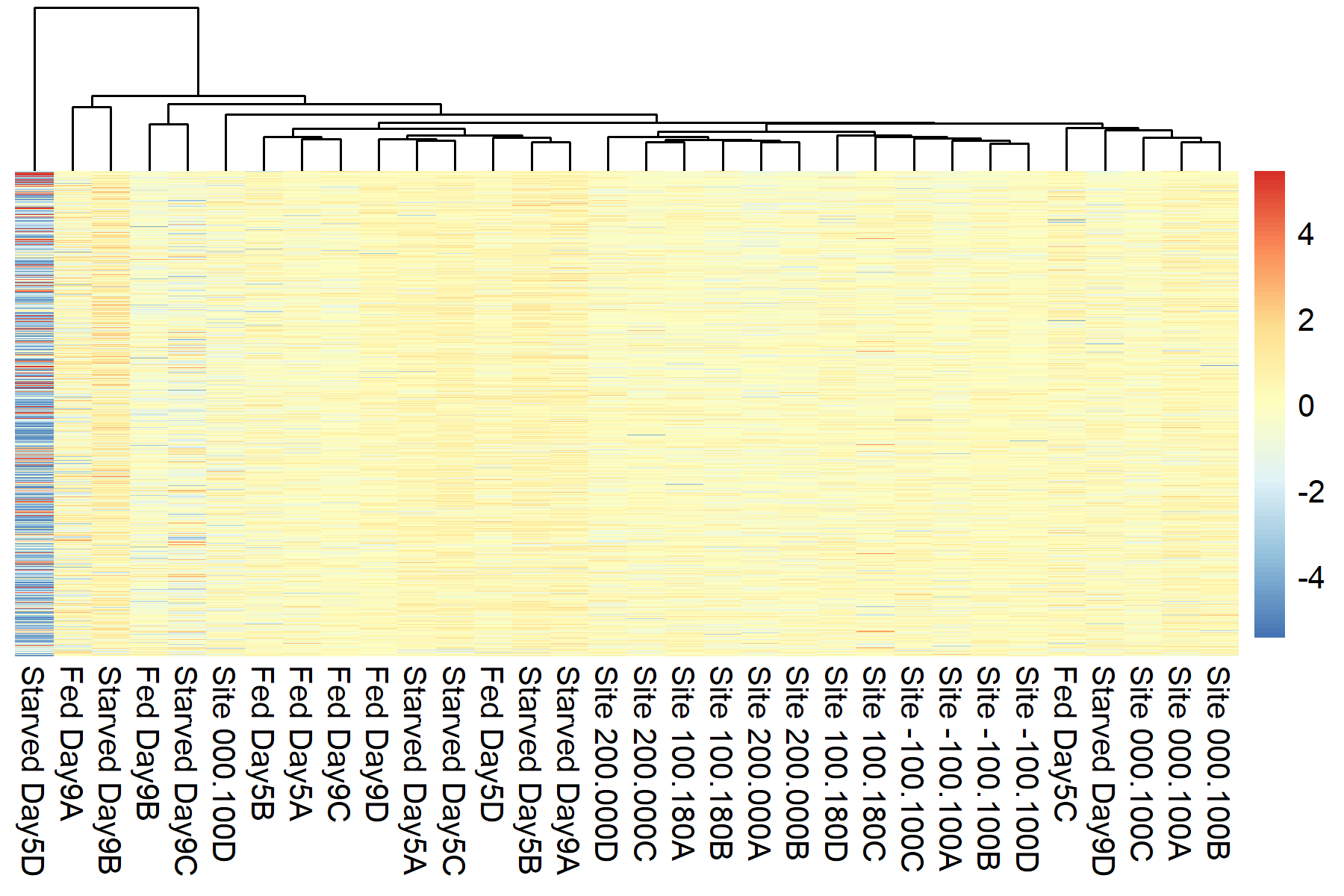

**Fig. S2.** Sample clustering based on sample distance matrix (A) and scaled and centered expression of the top 5000 most highly expressed genes (B) informed our decision to remove the outlier sample ‘Starved Day5D’ in *Calanus propinquus*.

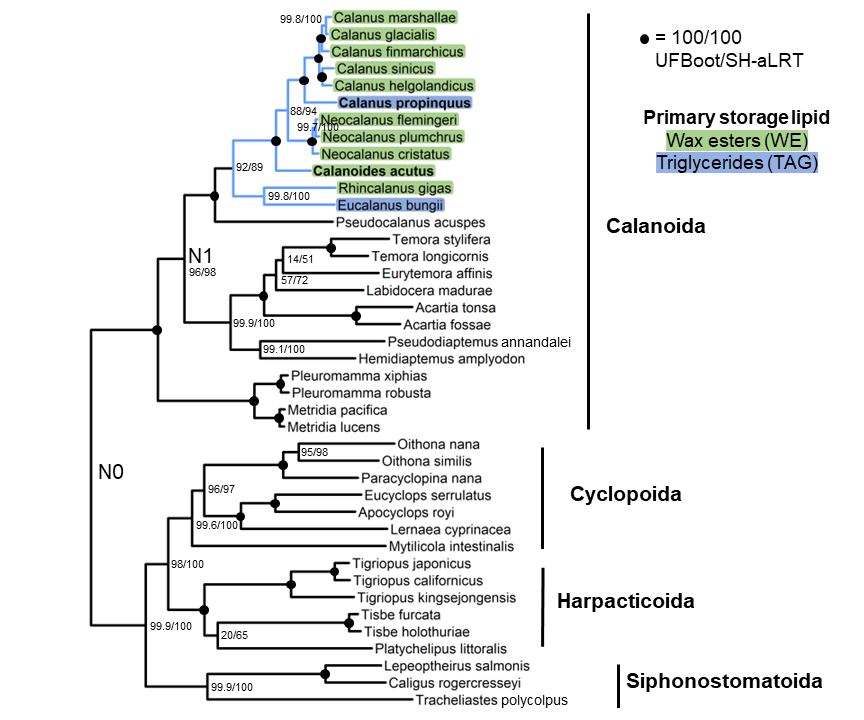

**Fig. S3.** Maximum-likelihood phylogeny constructed using edge-proportional partition model. Tree was constructed from 500 single-copy orthologs using IQ-TREE with the partitioning scheme and best model for each partition selected using PartitionFinder. Note that this tree has the same topology as Fig. 1, and was used as the guide tree for that analysis. Branch support values indicate ultra-fast bootstrap (UFBoot) and SH-aLRT, each calculated from 1000 replicates; black circles indicate 100% support from both metrics. Blue branches indicate the oil sac clade, and highlighted species names indicate primary storage lipid (WE or TAG). *Calanoides acutus* and *Calanus propinquus* are in bold. N0 indicates the root between Gymnoplea and Podoplea and N1 indicates the root of Calanoida.

**Food in gut	Reproductive content	Mortality**

***Calanoides acutus***

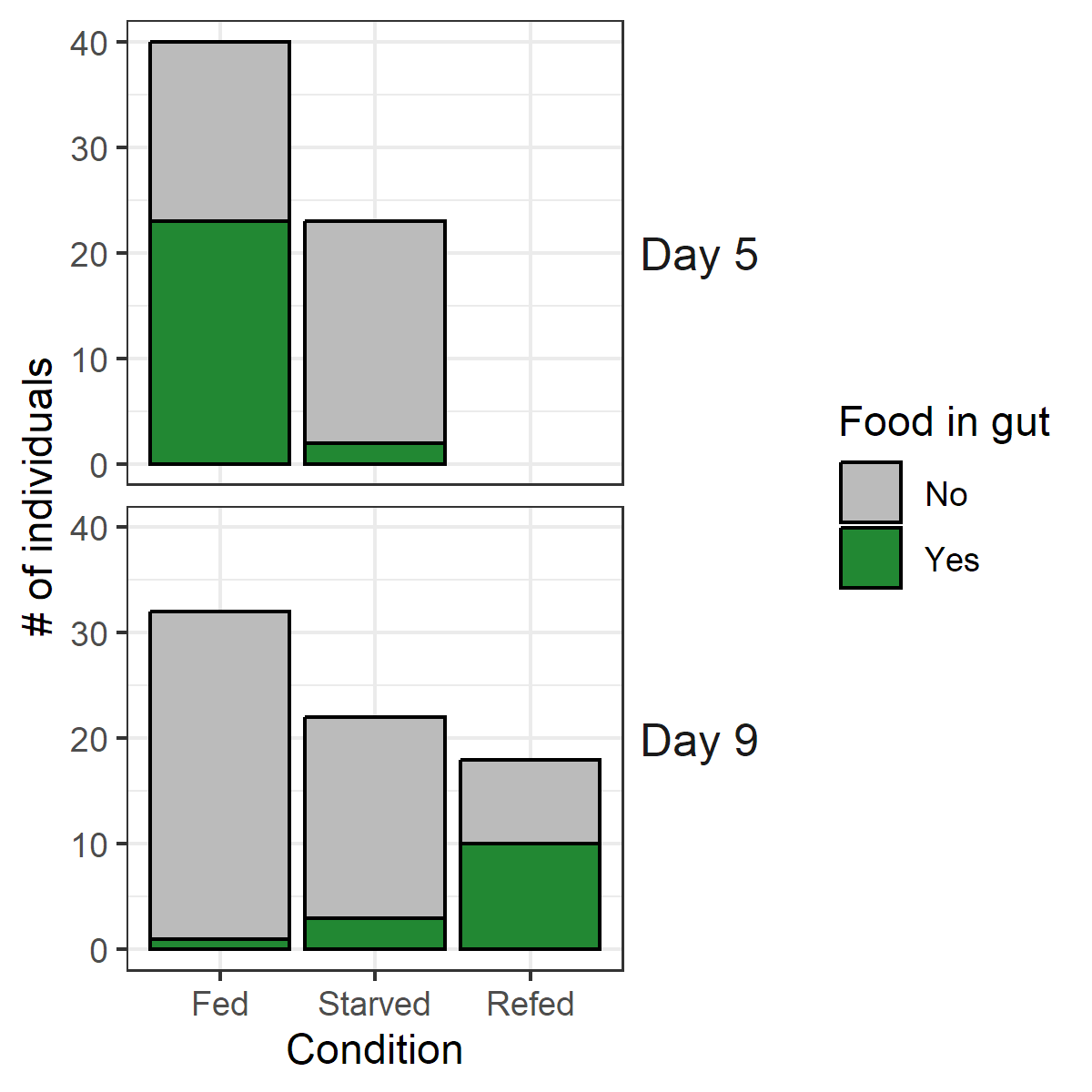

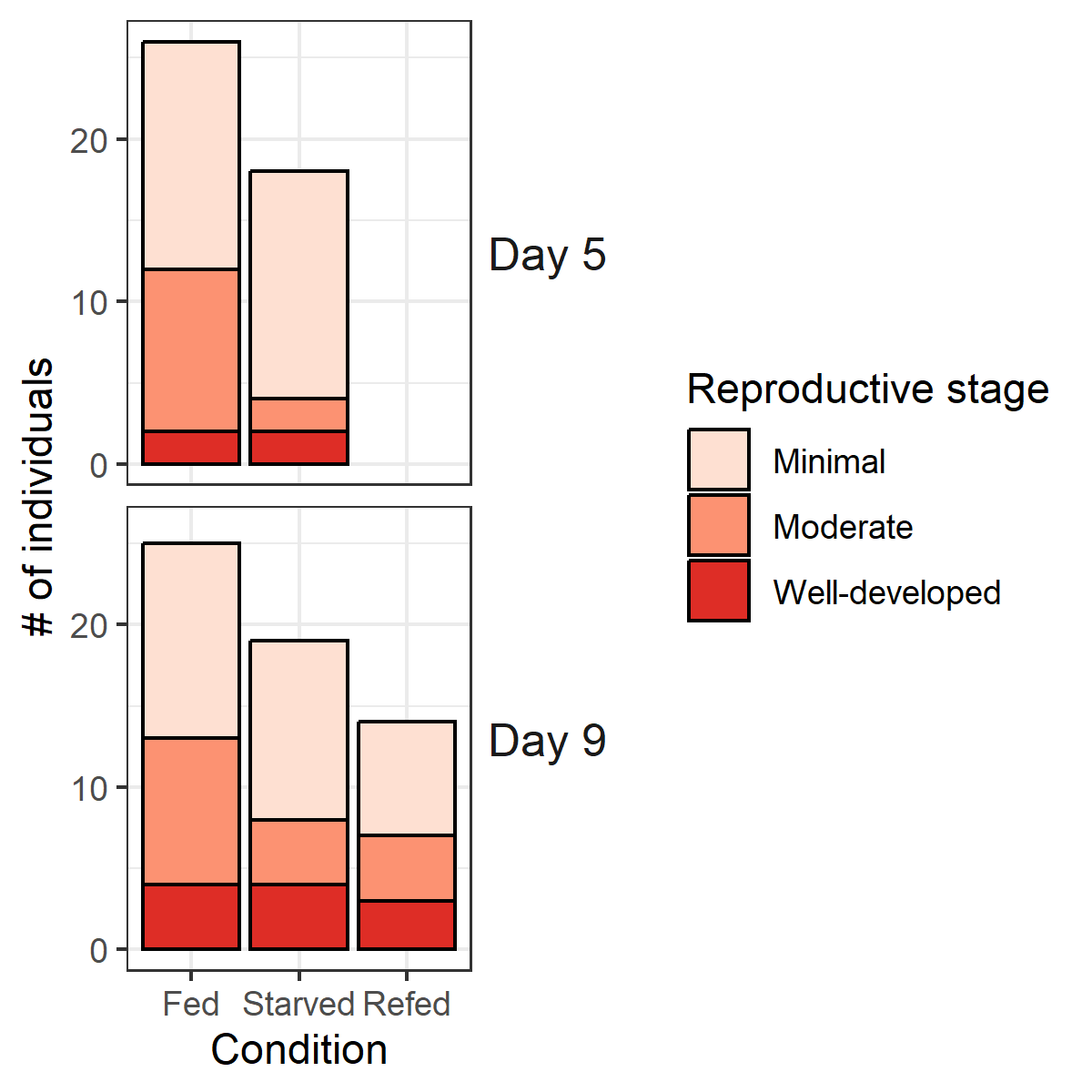

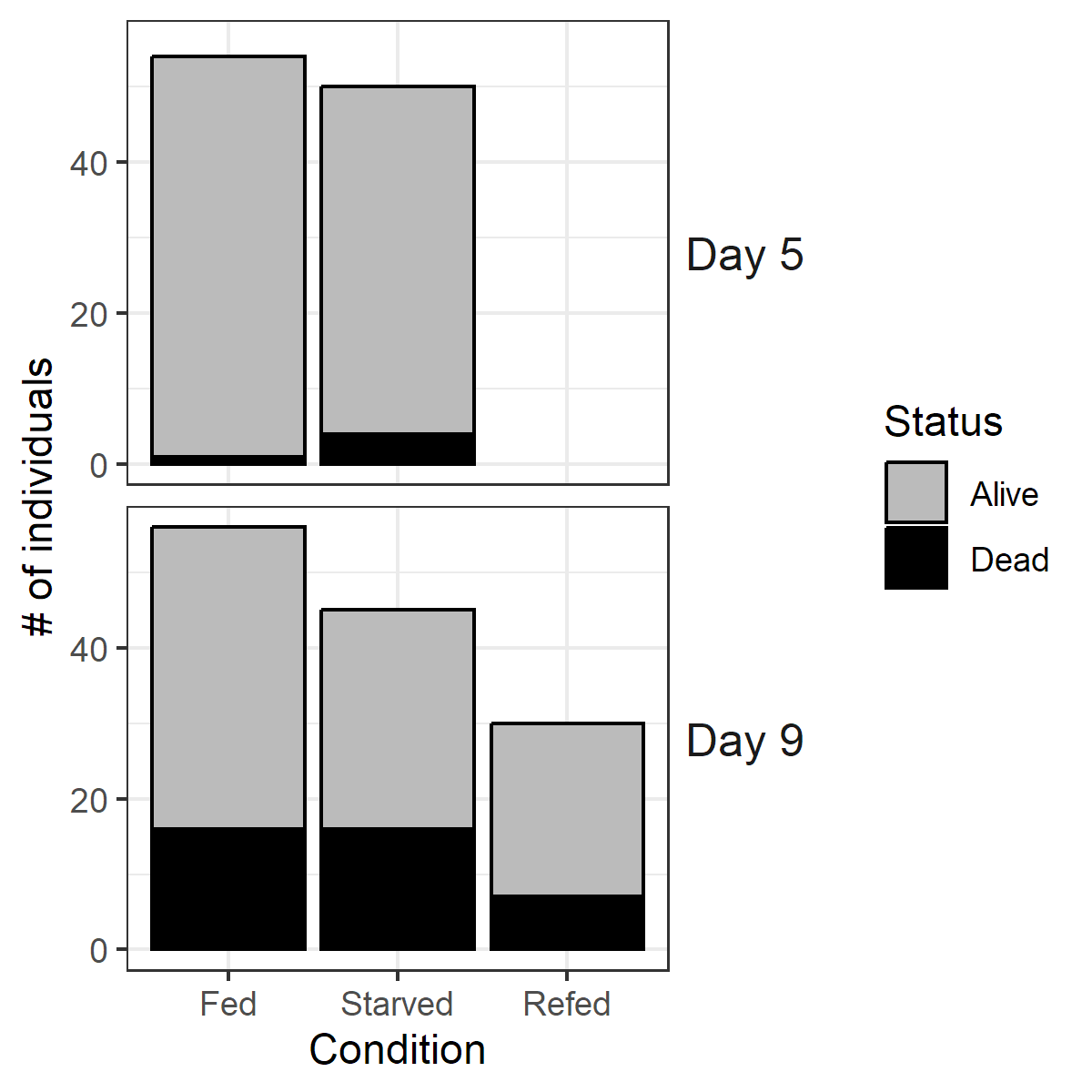

***Calanus propinquus***

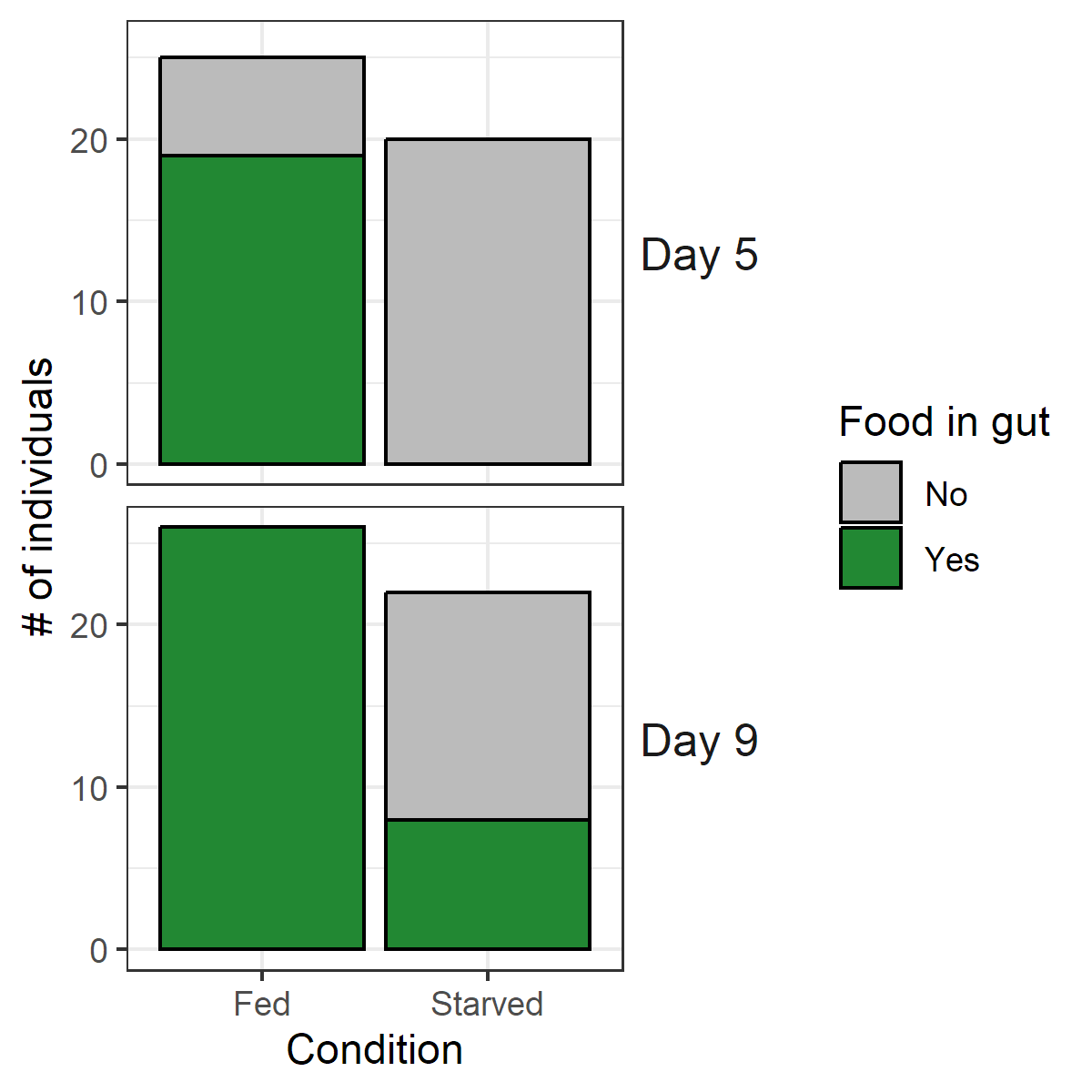

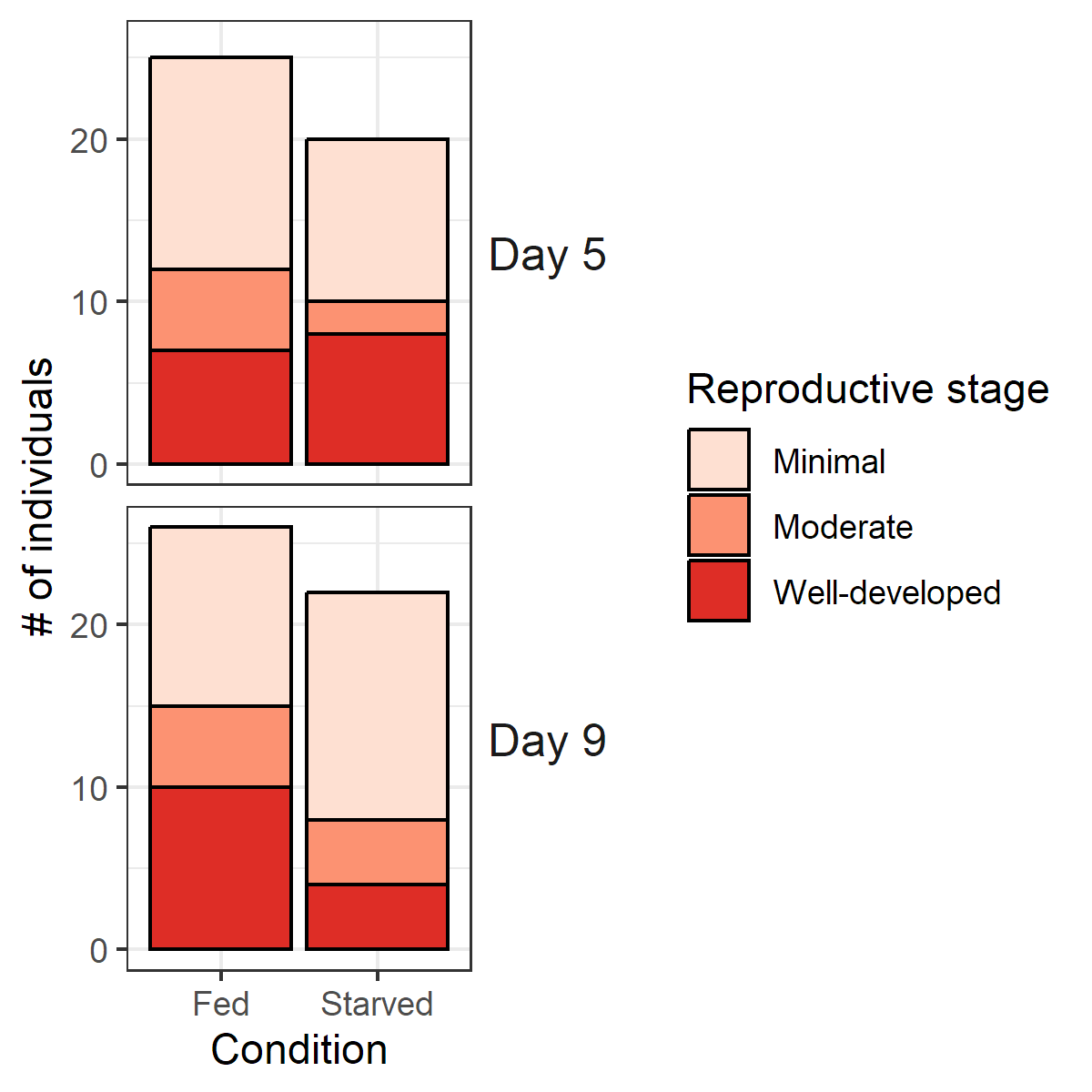

**Fig. S4.** Food in gut (left), reproductive condition (center), and mortality (right). Food in gut analysis is not quantitative (it does not contain information about the amount of food).

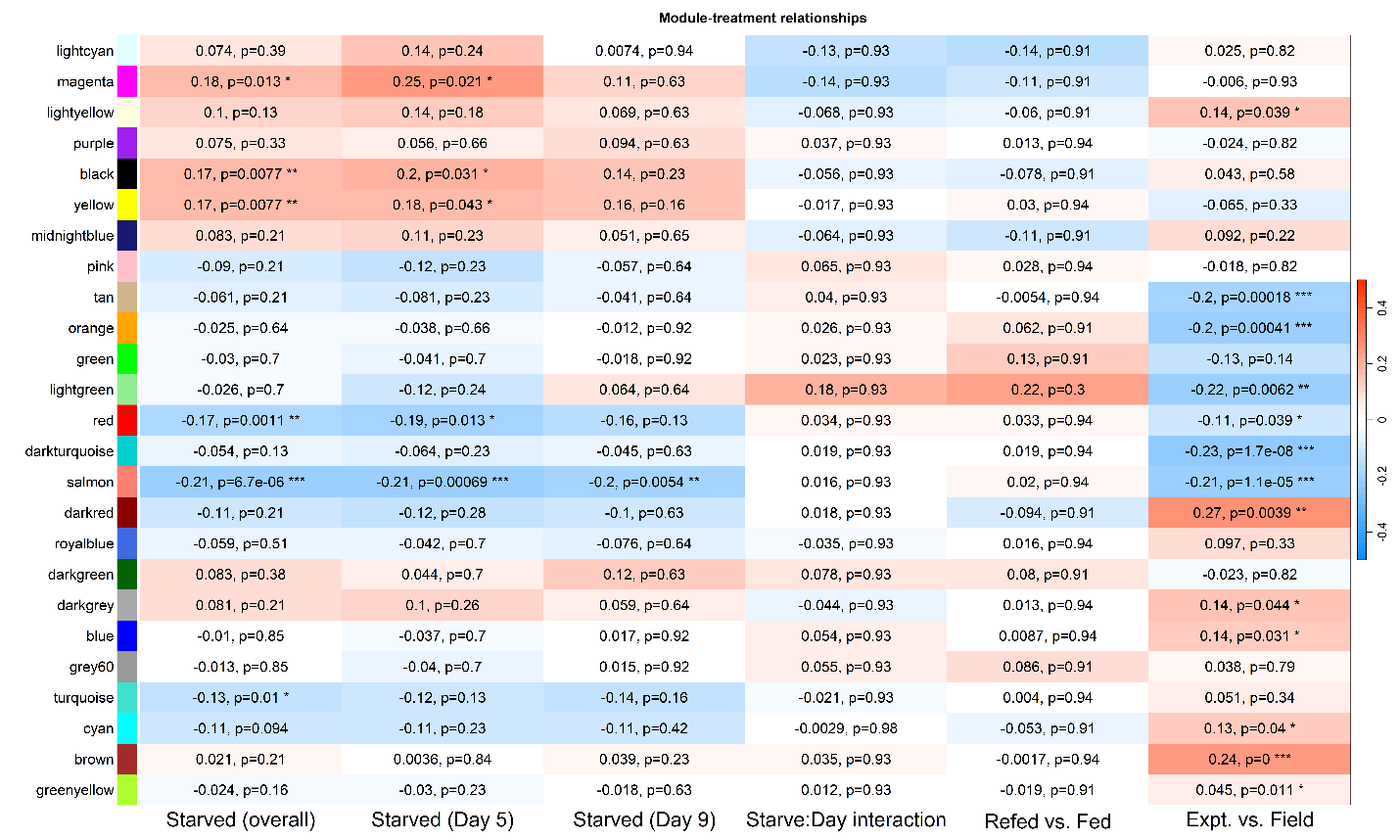

**Fig. S5.** Association analysis of *C. acutus* WGCNA modules with experimental variables. Associations were calculated using linear regressions of eigengenes, with the same model used for differential expression analysis. Cell colors represent model coefficients for a given contrast (mean change in eigengene expression); cells contain numeric coefficients and FDR-adjusted p-values (*, p < 0.05; **, p < 0.01; ***, p < 0.001).

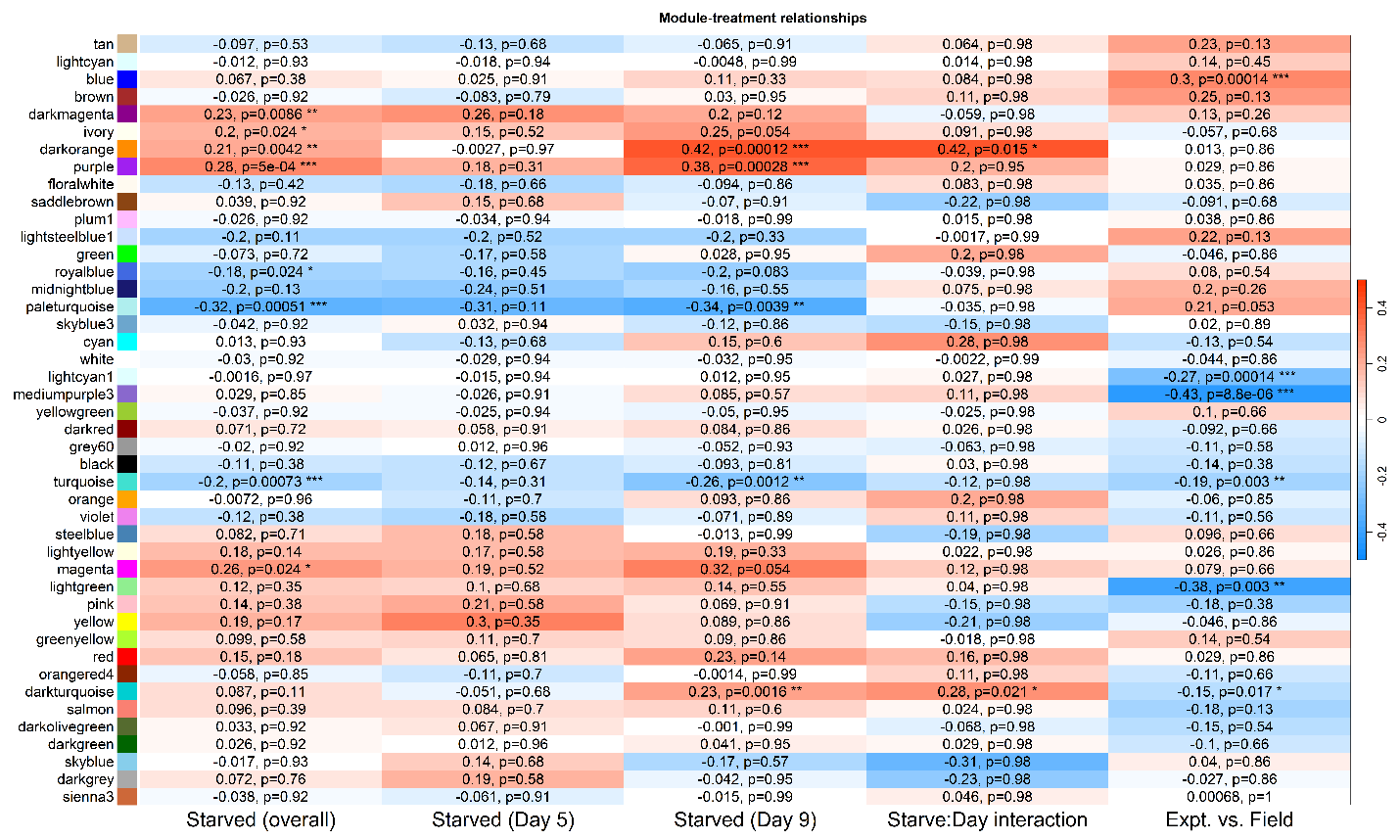

**Fig. S6.** Association analysis of *C. propinquus* WGCNA modules with experimental variables. Associations were calculated using linear regressions of eigengenes, with the same model used for differential expression analysis. Cell colors represent model coefficients for a given contrast (mean change in eigengene expression); cells contain numeric coefficients and FDR-adjusted p-values (*, p < 0.05; **, p < 0.01; ***, p < 0.001).

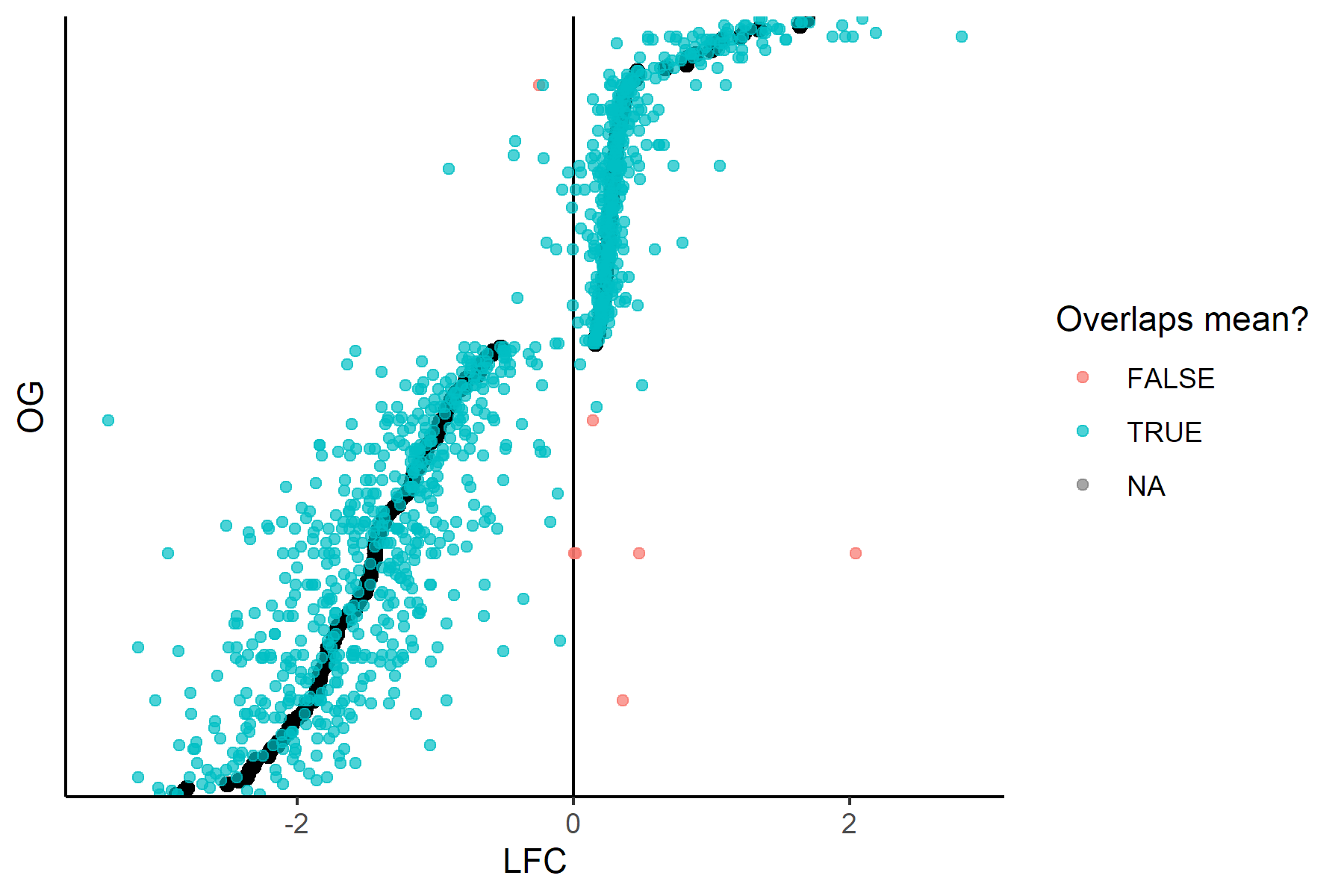

**Fig. S7.** Distribution of starvation log2 fold change (LFC) values in multi-copy conserved starvation-response genes (CSGs). Each black point represents the weighted mean LFC of an orthogroup, and each dot is the LFC point estimate for a gene. Red genes significantly differ from the mean (one-tailed t test, p<0.05) and have the opposite sign (7/1232 genes).

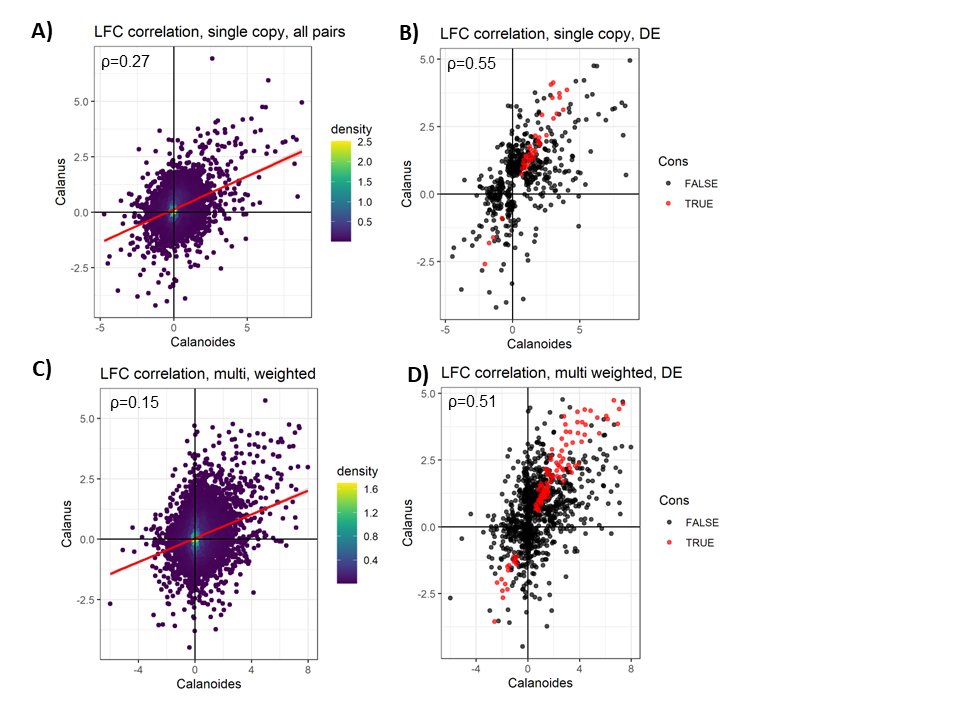

**Fig. S8.** Correlation of gene expression handling response between *Calanoides acutus* and *Calanus propinquus*. Left, correlation of handling LFC for single- (A) and multi-copy (C) genes. Right, LFC correlation for genes DE in at least one species for single- (B) and multi-copy (D) genes; conserved handling-response genes (CHGs) are in red. For multi-copy genes (bottom row), points represent the average value of the gene family weighted by the inverse squared standard error of each gene's estimate.

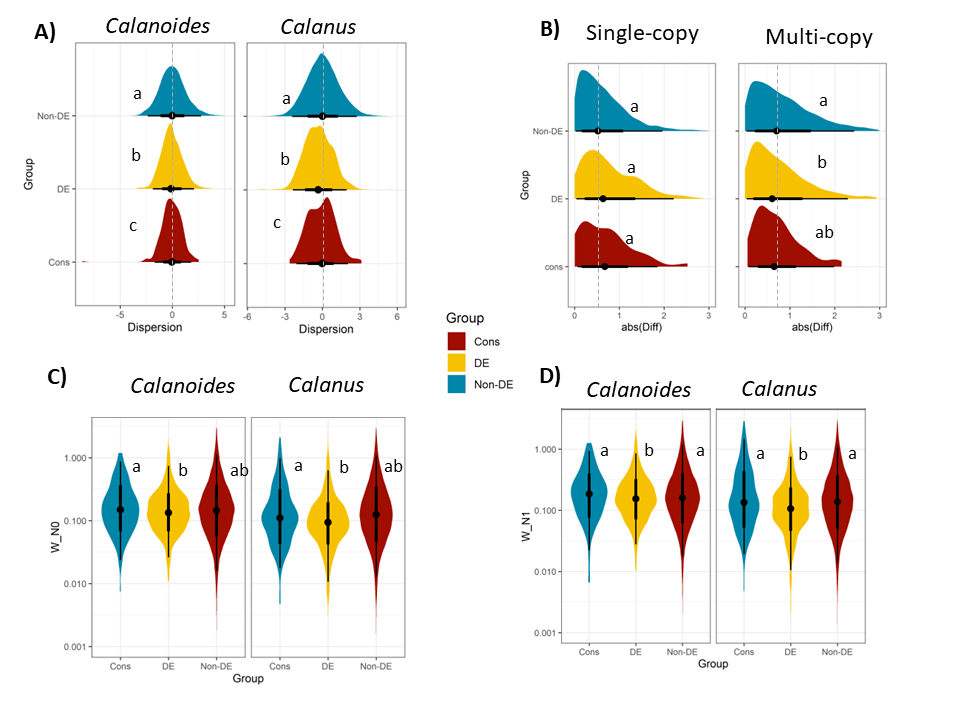

**Fig. S9.** Patterns of expression and sequence divergence for *handling-response* genes (HG). “Cons” refers to conserved handling-response genes (CHGs), and “DE” to species-specific handling-response genes. (A) Distribution of dispersion estimates for *Calanoides* *acutus* (left) and *Calanus propinquus* (right), calculated only using field samples. Letters indicate significant differences between groups in a weighted linear regression (LM, p<0.05), and dashed lines indicate non-DE group median. (B) Difference in dispersion estimates between species, for single-copy (left) and multi-copy (right) genes. For multi-copy genes, differences were calculated between average values of each gene family weighted by the inverse squared standard error of each gene's dispersion estimate. (C) Branch-wise dN/dS estimates in *Calanoides acutus* (left) and *Calanus propinquus* (right) calculated using gene trees inferred at the N0 node. Letters indicate significant differences between groups in a weighted logistic regression accounting for expression level (GLM, p<0.05). Note the log scale. (D) Branch-wise dN/dS estimates calculated using gene trees inferred at the N1 node.

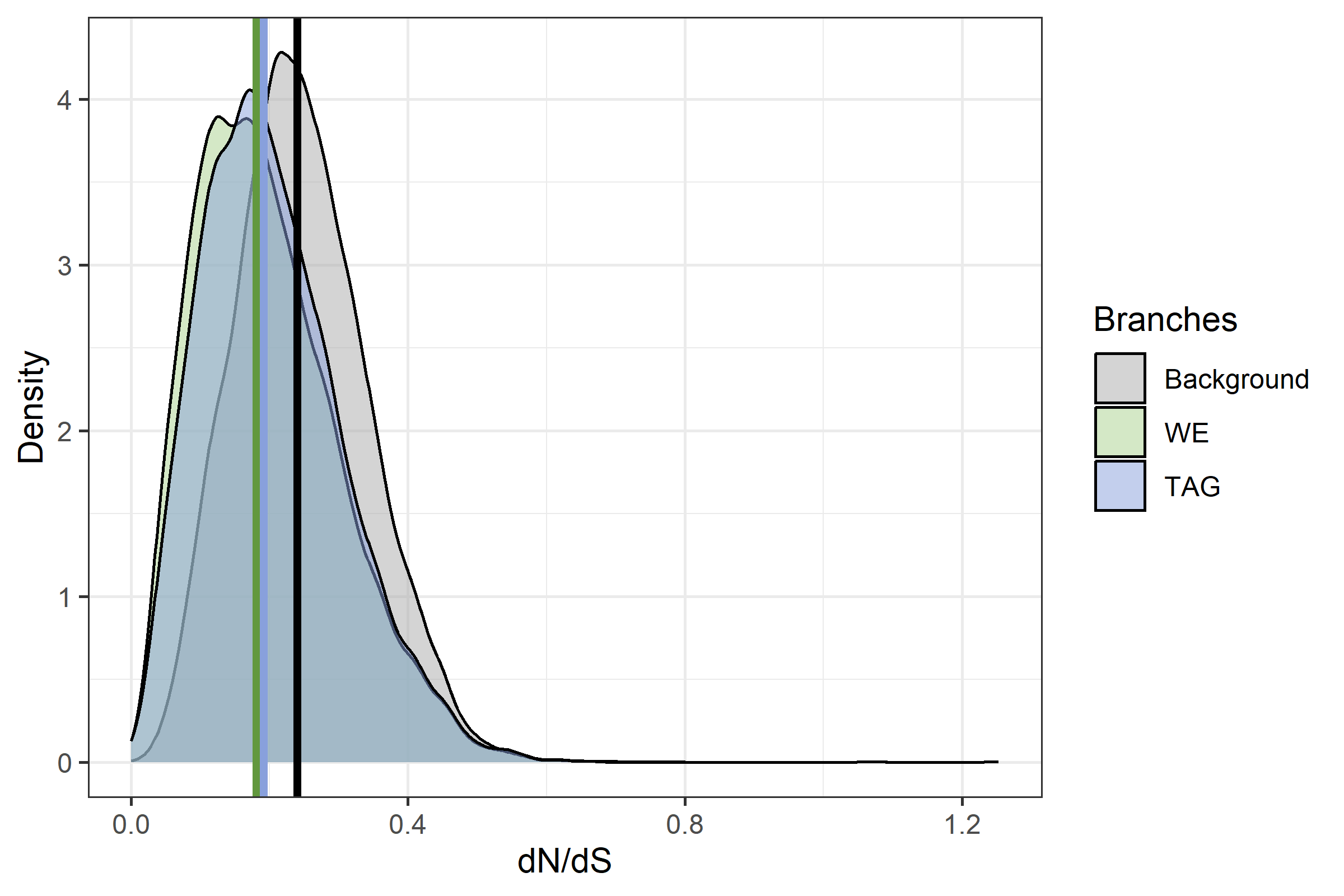

**Fig. S10.** Density plot of dN/dS estimates for background, WE, and TAG branches. These estimates were derived from codeml analysis of gene trees in the N1 dataset, averaged across one-ratio, two-ratio, and three-ratio genes. Vertical lines indicate median dN/dS for each group in gene trees.

A)	B)

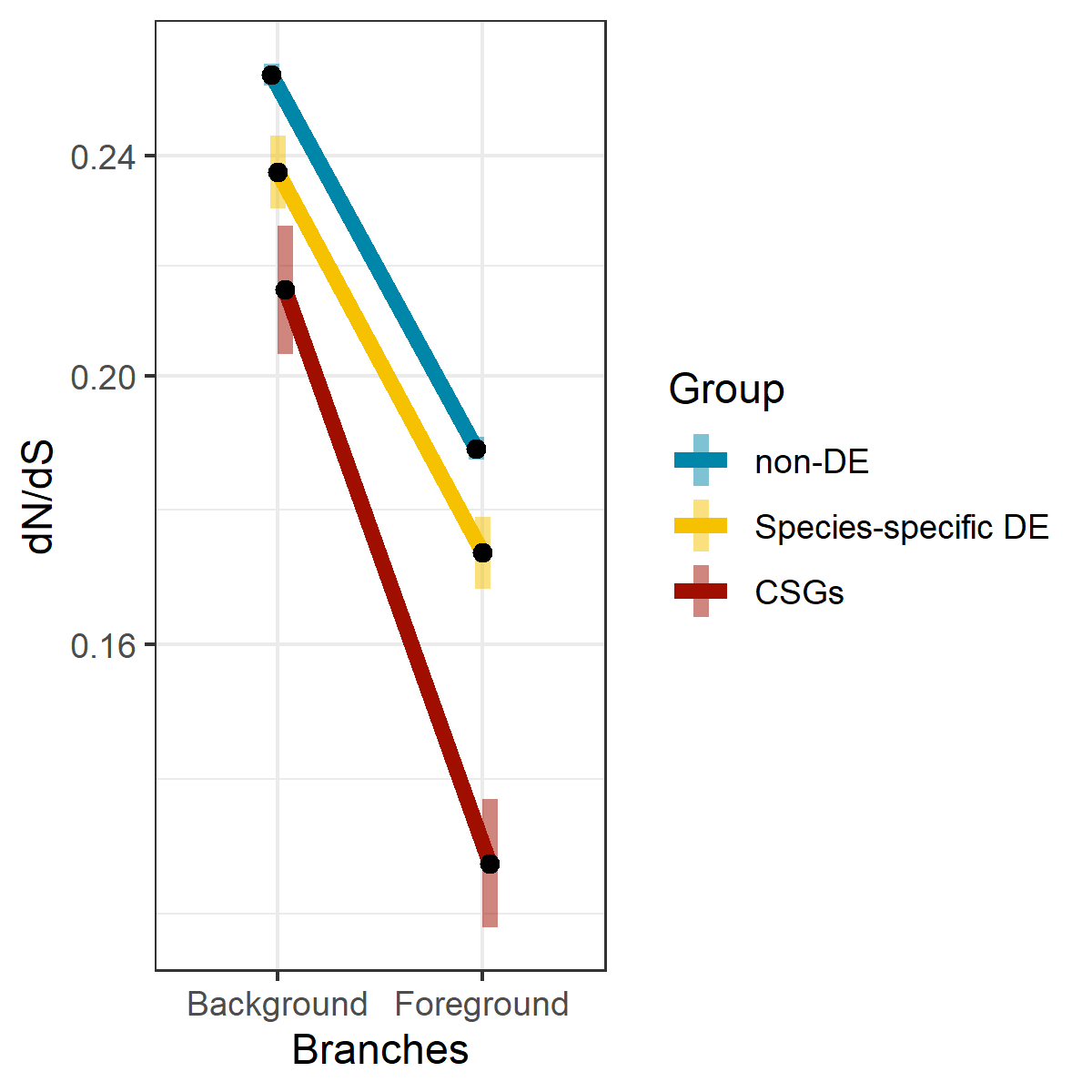

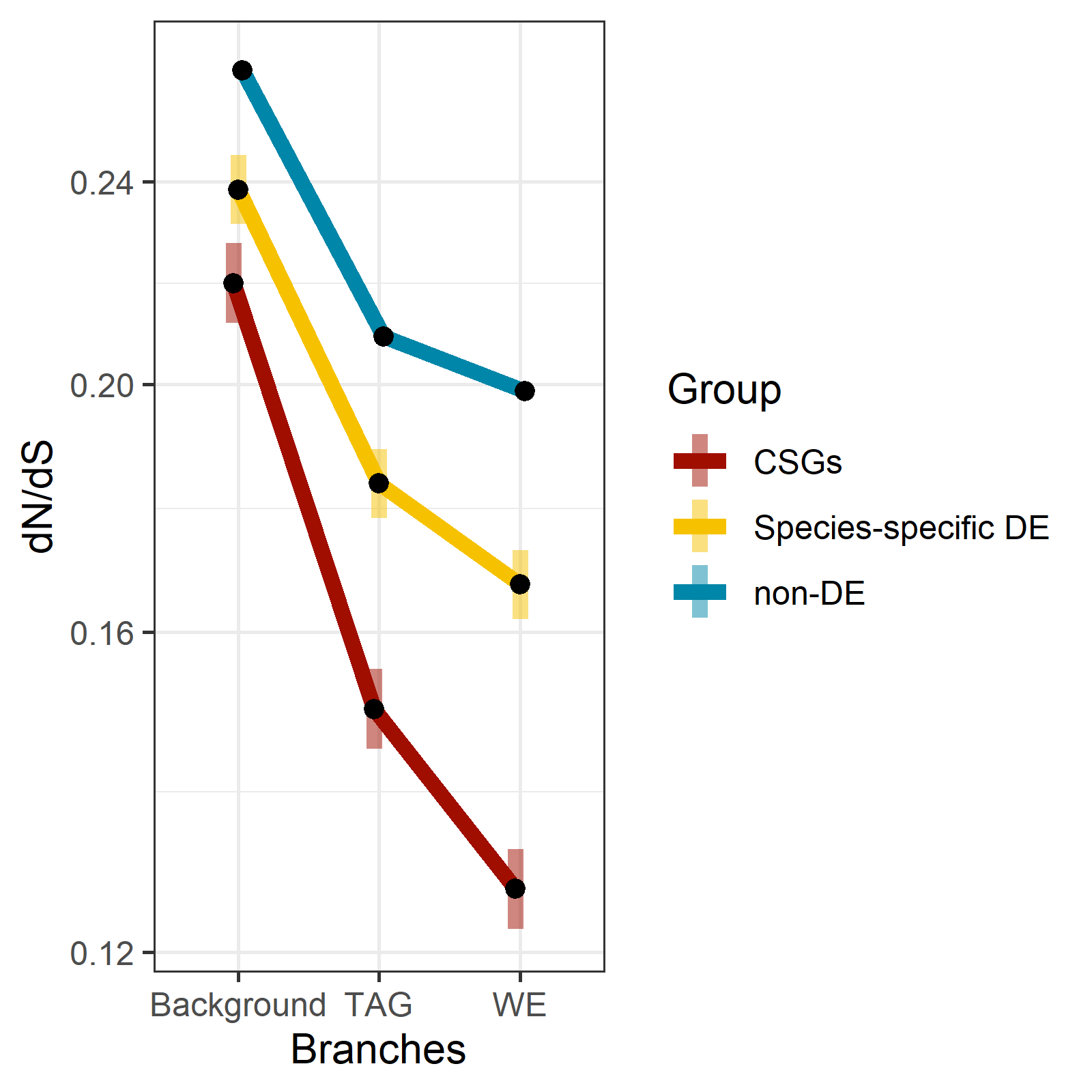
N0

C	D)

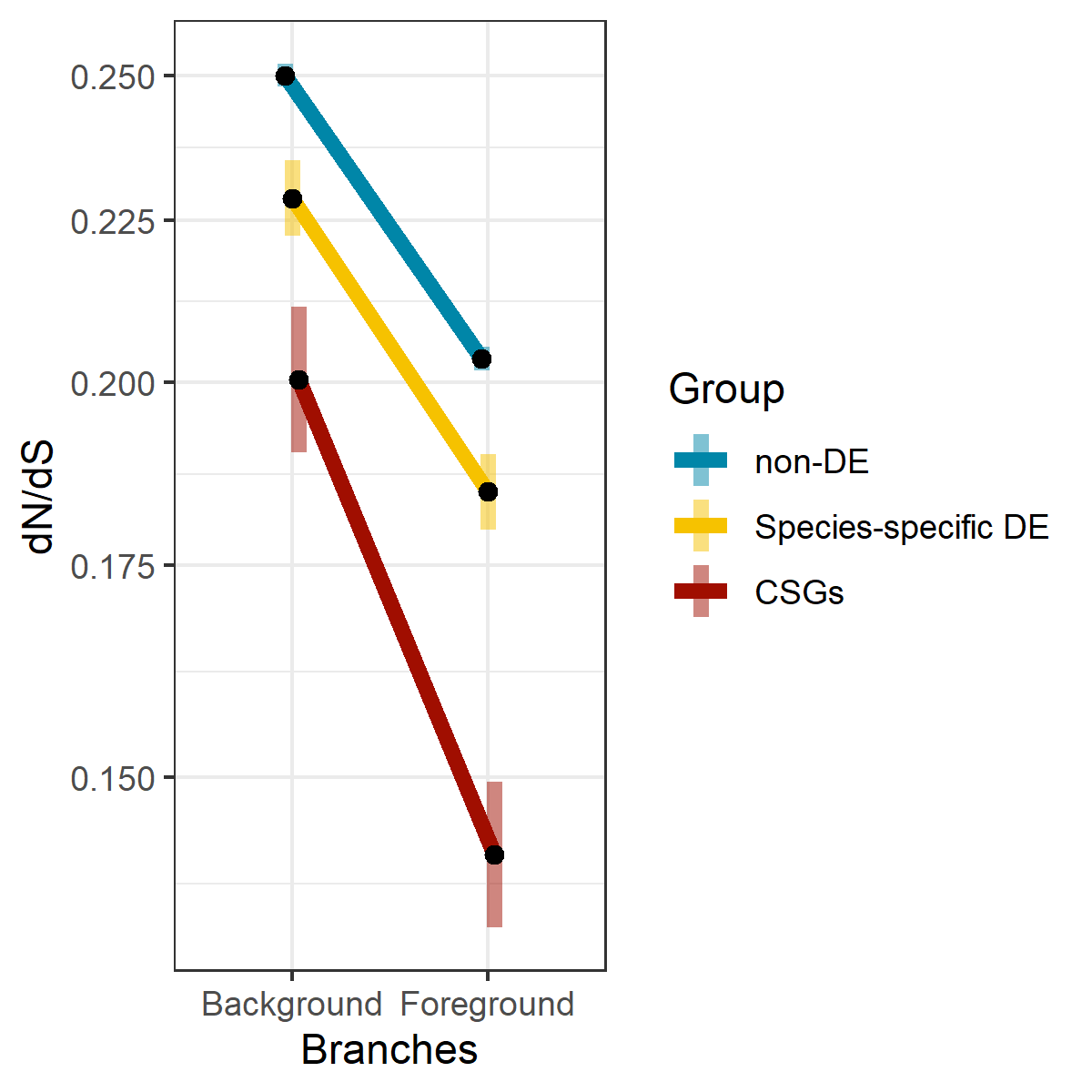

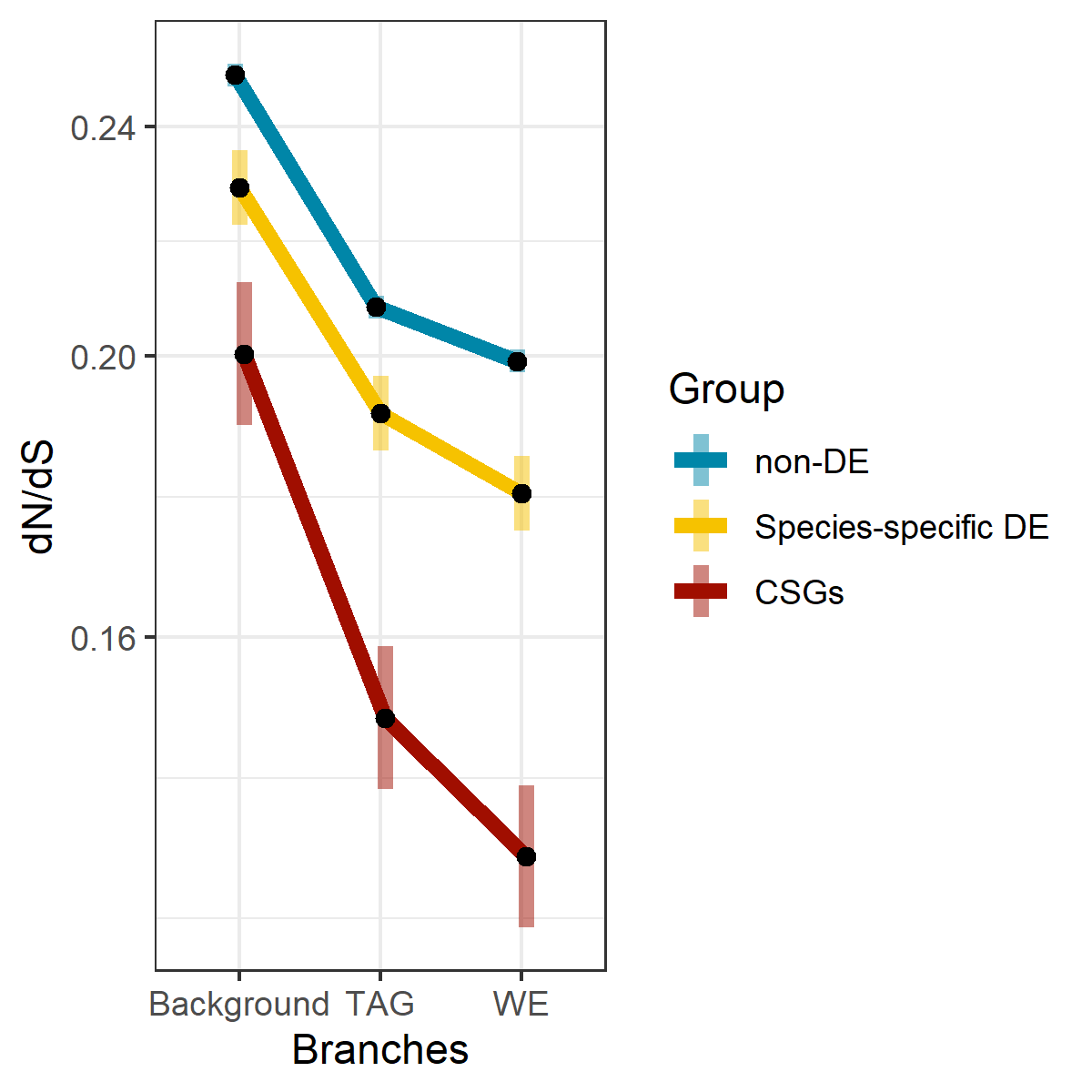
N1

**Fig. S11.** Interaction plots of estimated marginal means of codeml dN/dS estimates from log-link Gamma GLM for starvation-response genes. Left and right columns are identical, except right panels differentiate WE and TAG lineages within the foreground. A) and B) are derived from the N0 dataset (panel B is identical to Fig. 1C). C) and D) are derived from the N1 dataset. Shaded bars indicate 95% confidence intervals. Y-axis is on a natural log scale.

A)	B)

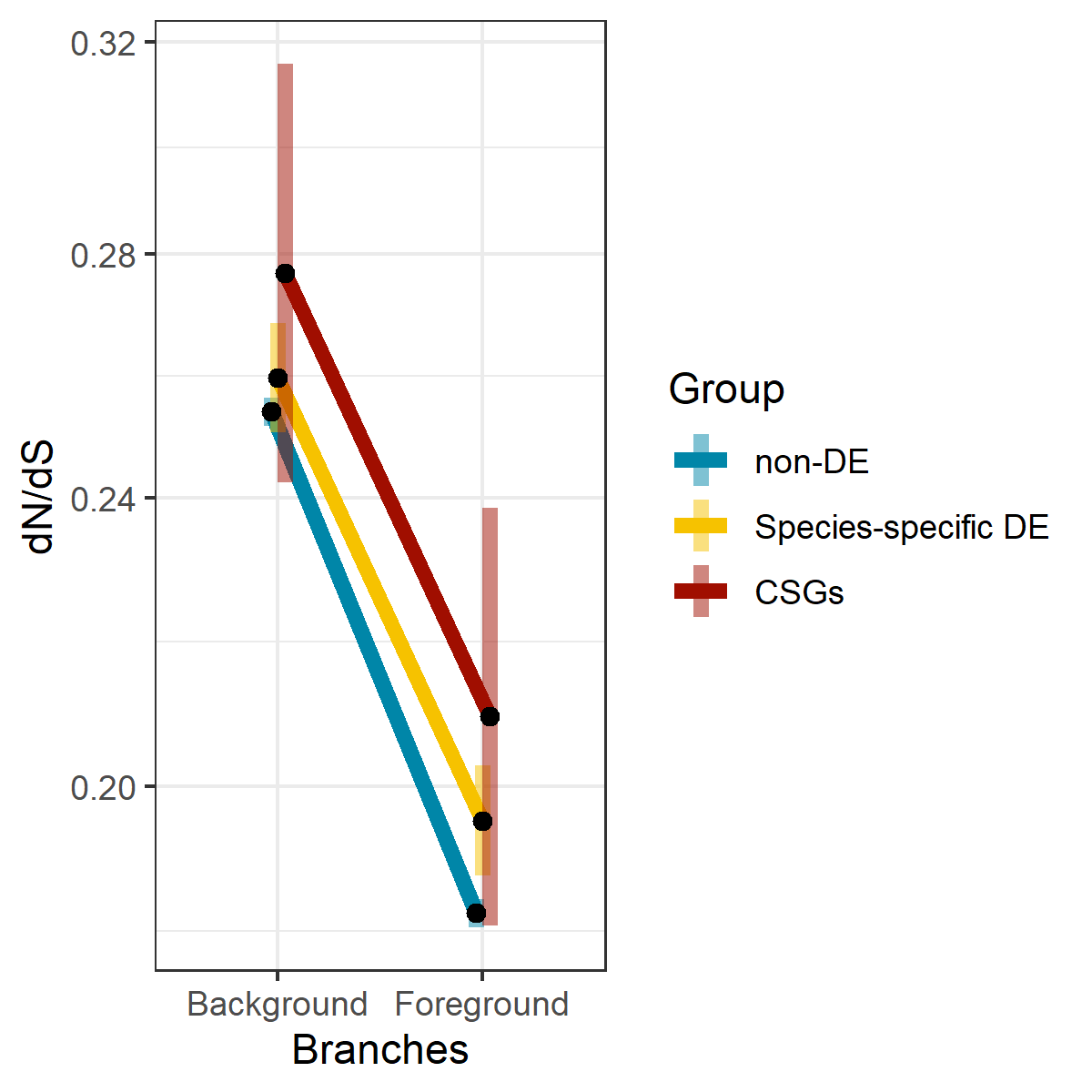

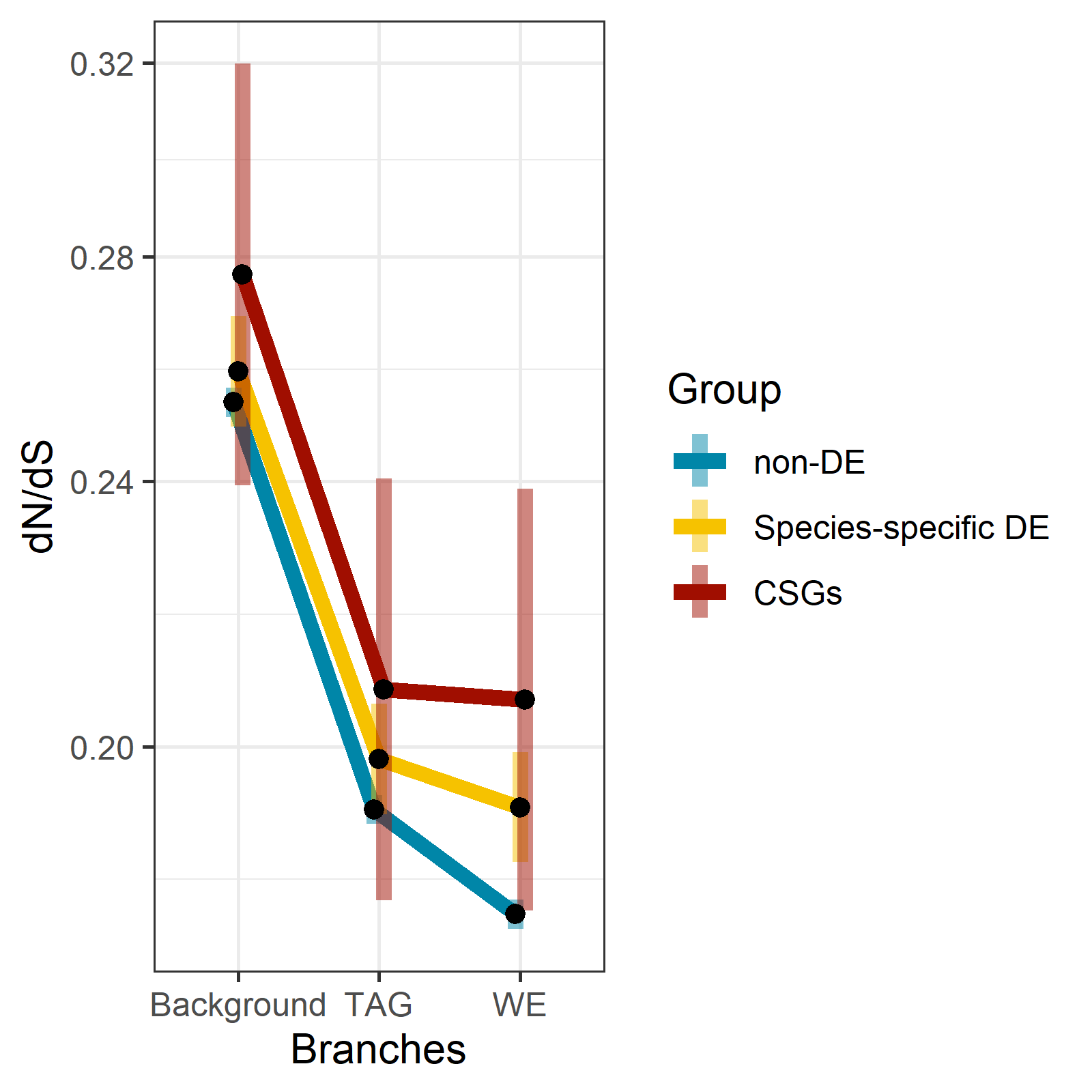
 N0

C)	D)

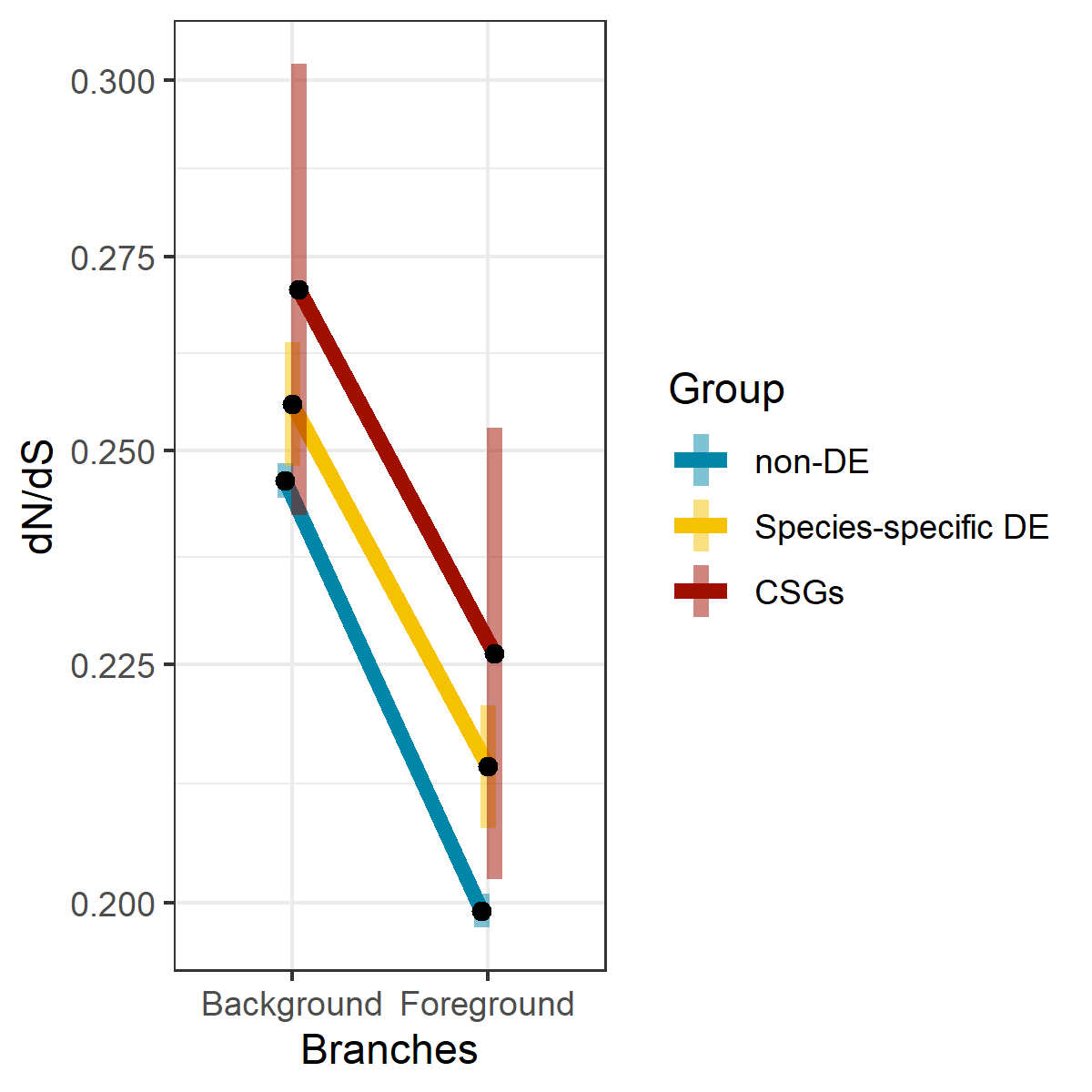

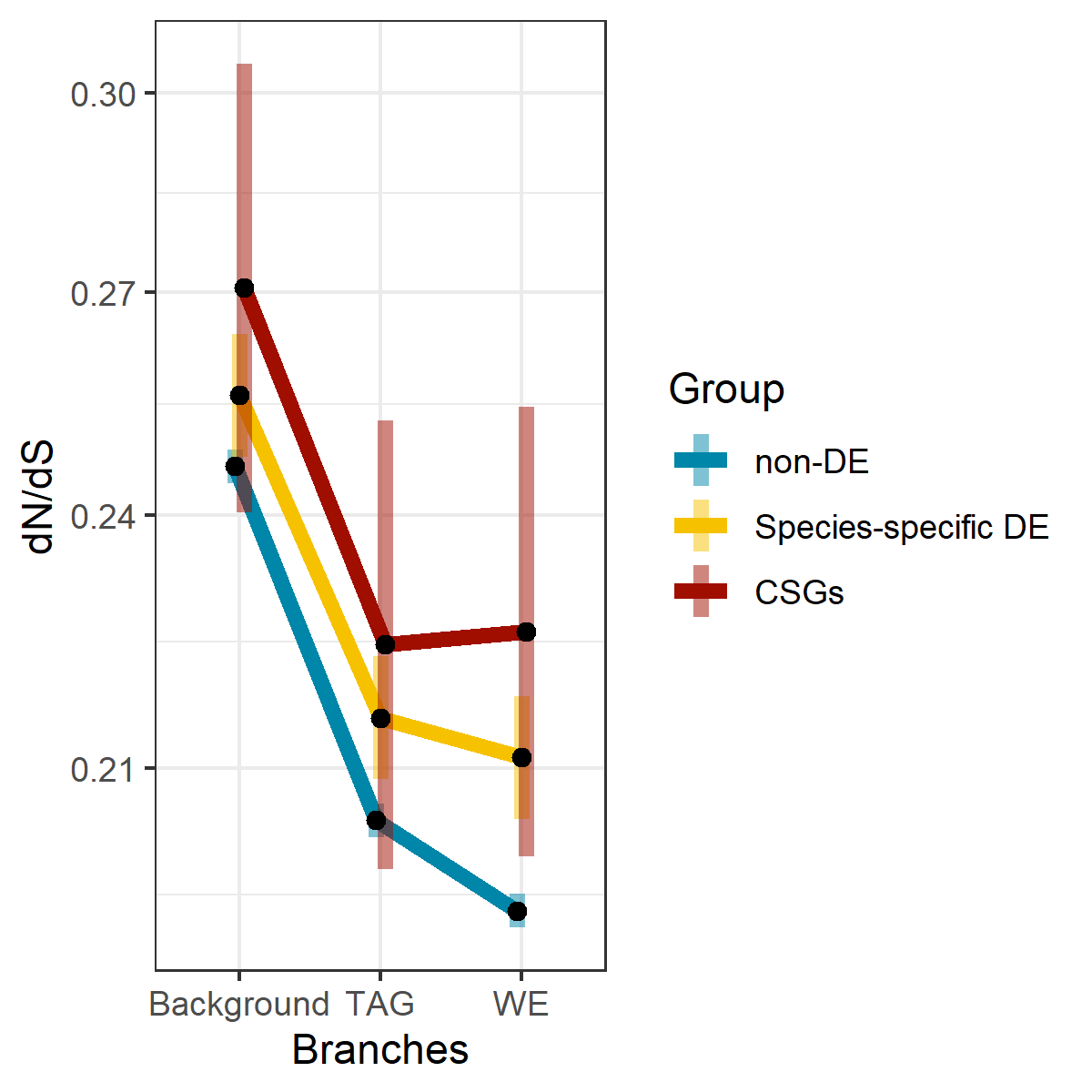
N1

**Fig. S12.** Interaction plots of estimated marginal means of codeml dN/dS estimates from log-link Gamma GLM for handling-response genes. Left and right columns are identical, except right panels differentiate WE and TAG lineages within the foreground. A) and B) are derived from the N0 dataset; C) and D) are derived from the N1 dataset. Shaded bars indicate 95% confidence intervals. Y-axis is on a natural log scale.
